# Transcriptional decomposition reveals active chromatin architectures and cell specific regulatory interactions

**DOI:** 10.1101/130070

**Authors:** Sarah Rennie, Maria Dalby, Lucas van Duin, Robin Andersson

## Abstract

Transcriptional regulation is tightly coupled with chromosomal positioning and three-dimensional chromatin architecture. However, it is unclear what proportion of transcriptional activity is reflecting such organisation, how much can be informed by RNA expression alone, and how this impacts disease. Here, we develop a transcriptional decomposition approach separating the proportion of expression associated with genome organisation from independent effects not directly related to genomic positioning.

We show that positionally attributable expression accounts for a considerable proportion of total levels and is highly informative of topological associating domain activities and organisation, revealing boundaries and chromatin compartments. Furthermore, expression data alone accurately predicts individual enhancer-promoter interactions, drawing features from expression strength, stabilities, insulation and distance. We further characterise commonalities and differences across predictions in 76 human cell types, observing extensive sharing of domains, yet highly cell-type specific enhancer-promoter interactions and strong enrichments in relevant trait-associated variants. Our work demonstrates a close relationship between transcription and chromatin architecture, presenting a novel strategy and an unprecedented resource for investigating regulatory organisations and interpretations of disease associated genetic variants across cell types.

## Introduction

The three dimensional organisation of a genome within a nucleus is strongly associated with cell-specific transcriptional activity (Gorkin et al., 2014). On a global level, transcriptional activation or repression is often accompanied by nuclear relocation of chromatin in a cell-type specific manner, forming chromatin compartments (Lieberman-Aiden et al., 2009) of coordinated gene transcription (Osborne et al., 2004; Schoenfelder et al., 2010) or silencing (Guelen et al., 2008). Locally, chromatin forms sub-mega base pair domains of self-contained chromatin proximity, commonly referred to as topologically associating domains (TADs) (Dixon et al., 2012; Nora et al., 2012). TADs frequently encompass interactions between regulatory elements, such as between promoters and enhancers (Heidari et al., 2014; Mifsud et al., 2015; Noordermeer et al., 2014; Rao et al., 2014) as well as between co-regulated genes (Nora et al., 2012), which reflects cell-type restricted transcriptional programs (Dixon et al., 2016). In contrast, ubiquitously expressed promoters are enriched close to domain boundaries (Dixon et al., 2012) and co-localise in the nucleus (Hug et al., 2017). A tight coupling between transcriptional activity and chromosomal positioning is further supported by positional clustering of co-expressed eukaryotic genes (Cohen et al., 2000; Michalak, 2008), a phenomenon that is preserved across taxa (Fukuoka et al., 2004). In addition, neighbouring gene expression correlation co-evolves and is particularly evident at distances below a mega base pair (Ghanbarian and Hurst, 2015). These observations are in line with coordinated gene expression within TADs (Le Dily et al., 2014; Nora et al., 2012), suggesting that the coupling between gene expression and chromatin architecture is, at least partly, linked to chromosomal positioning.

Genetic disruption or chromosomal rearrangements of TAD boundaries may result in aberrant regulatory transcriptional activities (Krijger and de Laat, 2016). While this suggests a key role for chromatin structure in correct transcriptional activity, a general directional cause and effect between chromatin architecture and transcriptional activity is unclear. On one hand, TAD boundaries are to a large extent invariant between cell types (Dixon et al., 2012; Nora et al., 2012) and have been suggested to be mainly formed by architectural proteins (de Wit et al., 2015; Nora et al., 2017; Phillips-Cremins et al., 2013; Rao et al., 2014) independent of transcription (Hug et al., 2017). On the other hand, transcription seems to play a major role in the maintenance of regulatory organisation within TADs (Hug et al., 2017; Li et al., 2015). In addition, chromatin compartments are not affected by architectural protein depletion (Nora et al., 2017). Rather, nuclear three-dimensional co-localisation of genes seems to be driven to a large degree by transcriptional activity (Branco and Pombo, 2006; Hug et al., 2017; Mitchell and Fraser, 2008; Ulianov et al., 2016).

The strong relationships between transcription, chromosomal positioning and three-dimensional chromatin organisation can be exploited for inference of one feature from another. Compartments of transcriptional activity can be deduced from genome-wide chromatin interaction data (Lieberman-Aiden et al., 2009; Rao et al., 2014) and, inversely, co-expression is indicative of enhancer-promoter (EP) interactions (Andersson et al., 2014). In addition, RNA expression was placed among the top ranked features for predicting EP interactions in a recent machine learning approach (Whalen et al., 2016). However, it is unclear what proportion of expression is associated with chromatin topology and chromosomal positioning and what proportion is reflecting regulatory programs independent of the former. A way to systematically extract and separate these components from expression data could lead to new insights into chromatin topology and cell-type specific transcriptional regulation. This is of high importance since genome-wide profiling of chromatin interactions using chromatin conformation capture techniques such as HiC (de Wit and de Laat, 2012) are largely intractable due to several limitations including low spatial resolution, high cost and requirements of large abundant cellular material.

In this study, we establish a transcriptional decomposition approach to investigate and model the coupling between transcriptional activity, chromosomal positioning and chromatin architecture. Via modelling of expression similarities between neighbouring genomic loci, we show that the proportion of expression related to genome architecture can be separated from effects independent of genomic positioning, and that both components are strongly represented, with contributions depending on cell type and location. We demonstrate that the positionally dependent component is highly reflective of chromatin organisation, revealing chromatin compartments and structures of transcriptionally active TADs. We further demonstrate how transcriptional components can be used to infer cell type-specific chromatin interactions and find informative transcriptional features, including enhancer expression strength, that distinguish long-distance interacting from non-interacting EP pairs. We demonstrate the accuracy of our approach in well-established cell lines and then decompose expression data from 76 human cell types in order to investigate their active chromatin architectures. Our results indicate extensive sharing of expression-associated domain structures across human cells, reminiscent of that observed for active TADs. Promoter-localised, positionally independent events as well as EP interactions were, on the other hand, found to be highly cell-type specific. Lastly, we demonstrate how transcriptional components and predicted EP interactions can be used to gain insights into the genetics consequences of complex diseases. We observe variable enrichments of genetic variants in expression components across cell types and traits and find several EP interactions in which disease-associated SNPs at enhancers may cause aberrant expression of distal genes. Taken together, we observe a tight coupling between transcriptional activity and three-dimensional chromatin architecture and suggest that regulatory topologies, domain structures, and their activities may be inferred by RNA expression data and chromosomal position alone.

## Results

### Decomposition of expression data reveals chromatin compartments and independent gene regulatory programs

We posited that a proportion of steady state RNA expression is reflecting three-dimensional chromatin organisation (Fig. 1A). We reasoned that a transcription unit (TU) is likely to be more similar in terms of expression to its linearly proximal TUs than to distal loci, which are likely to be associated with different domains of chromatin interactions. However, expression is influenced by gene-specific transcriptional and post-transcriptional regulatory programs independent of a TU’s chromosomal position. Thus, in order to investigate the coupling between transcriptional activity and genome organisation, we needed to be able to estimate the proportion of expression from a genomic region that can be explained by its chromosomal position. To this end, we developed a transcriptional decomposition approach (Fig. 1B) to separate the component of expression reflecting an underlying positional relationship between neighbouring genomic regions (positionally dependent (PD) component) from the expression dictated by TUs’ individual regulatory programs (positionally independent (PI) component). Our strategy is based on approximate Bayesian modelling (Rue et al., 2009) and utilises replicate measurements to decompose normalised aggregated RNA expression read counts in genomic bins into two components (PD, PI) and some constant intercept (see Methods).

**Figure 1.**
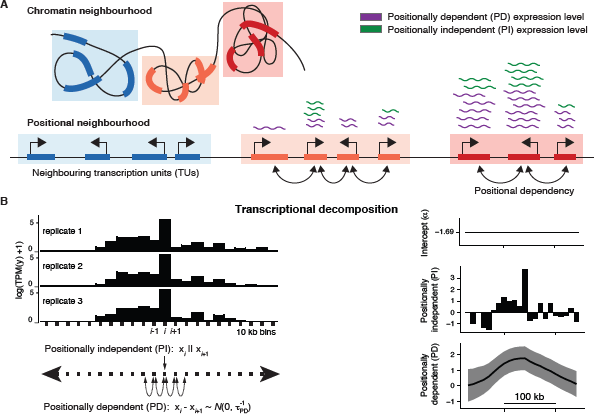
Transcriptional decomposition separates the proportion of RNA expression related to chromosomal position from positionally independent effects. **A**: Schematic illustrating how RNA expression derives from two major sources. The positionally dependent (PD) component reflects the underlying dependency between linearly proximal TUs in chromosomal, positional neighbourhoods, which are related to chromatin neighbourhoods of TU three-dimensional proximity. The positionally independent (PI) component reflects localised, gene-specific regulatory programs, unaffected by the positioning of TUs. **B**: Overall strategy of how replicated samples are decomposed into transcriptional components. Via approximate Bayesian modelling, normalised RNA expression count data quantified in genomic bins (here 10kb), are decomposed into an intercept (*α*), a PI component and a PD component. The PD component is modelled as a first-order random walk, in which the difference between consecutive bins is assumed to be Normal and centred at 0 (see Methods).

We applied transcriptional decomposition to replicated Cap Analysis of Gene Expression (CAGE) (Kanamori-Katayama et al., 2011) data (FANTOM Consortium and the RIKEN PMI and CLST (DGT) et al., 2014), measuring transcription initiation sites and steady-state abundances of capped RNAs, from GM12878, HeLa-S3, and HepG2 cells. In the modelling, we used 10kb genomic bins to capture large-scale positional effects. We first asked whether the information content of the two components contained different biological interpretations as posited. Upon close inspection (Fig. 2A), the PD component displayed considerably broader patterns than the PI component and appeared highly similar between cell types. Despite overall similarities in the PD component, we identified large differences in individual loci between cells, as exemplified by *KCNN3* and *NOS1AP* genes (Fig. 2B,C). For instance *NOS1AP* is in GM12878 cells associated with low PD signal and appears to reside in polycomb-repressed chromatin, as indicated by high levels of histone modification H3K27me3 and low levels of histone modification H3K36me3. HepG2 and HeLa-S3 cells displayed opposing signals for this locus, suggesting that the PD component contains information about chromatin compartments.

**Figure 2.**
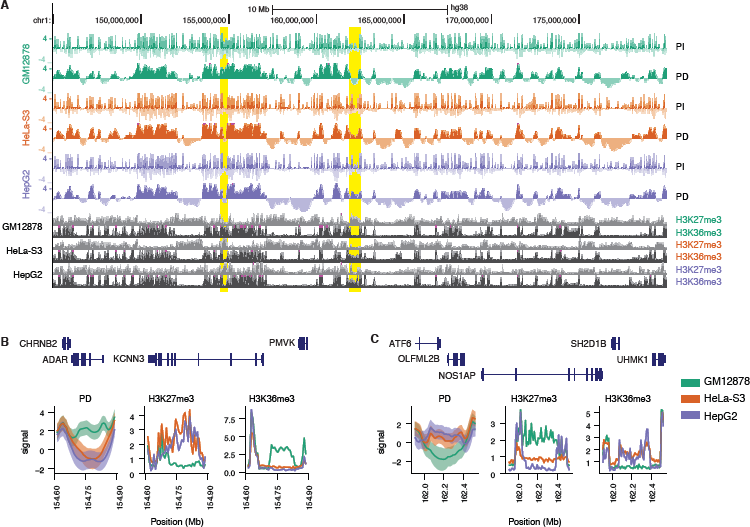
Transcriptional decomposition across chromosomes. **A**: PI and PD components, as well as H3K27me3 and H3K36me3 ChIP-seq data for GM12878, HeLa-S3 and HepG2 cells at locus chr1:145,000,000-180,000,000. **B-C**: Loci (highlighted in A) around *KCNN3* (**B**) and *NOS1AP* (**C**) genes showing cell-type specific PD signals. The PD signal and ChIP-seq data associated with repression (H3K27me3) and activation (H3K36me3) are shown.

In order to understand the observed effects on a genome-wide scale, we compared the PD component with HiC-predicted chromatin compartments (Rao et al., 2014) in GM12878 cells. We observed that the PD signal in regions of active chromatin was higher than in those of facultative or constitutive chromatin, while constitutive chromatin states had the lowest signal (Fig. 3A). Overlaying the PD component on HiC compartment boundaries also showed clear shifts, many magnitudes stronger than what could be detected using the PI component (Supplementary Fig. S1A). In addition, we observed that the PD component clearly correlated within compartments more strongly than between compartments (Supplementary Fig. S1B-E). These results show that the states and boundaries of compartments are reflected by positionally attributable expression (PD signal) and its relative change between consecutive bins.

**Figure 3.**
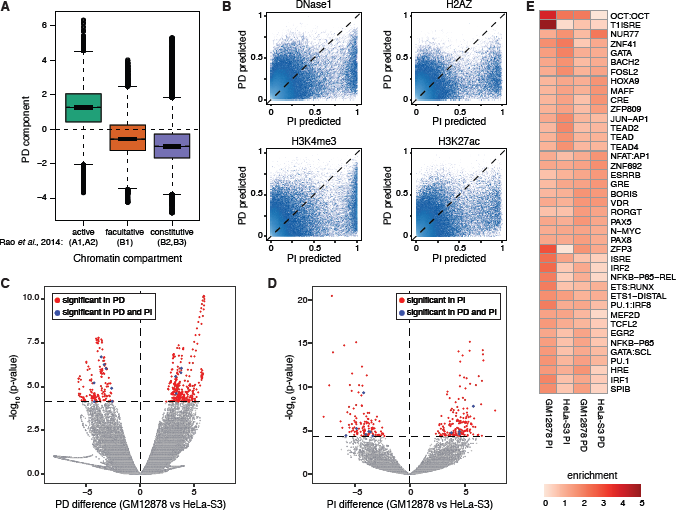
Transcriptional components reveal chromatin compartments and localised promoter-associated effects. **A:** Box-and-whisker plot of GM12878 PD signal grouped according to HiC-derived chromatin compartments. **B:** Random forest class (presence/absence) probability of DNase1, H2AZ, H3K4me3, and H3K27ac as learned from the PI component (x axes) and the PD component (y axes). **C:** PD difference (x axis) versus FDR-adjusted p-value (rescaled by -log_10_) for PD component differential expression between GM12878 and HeLa-S3. Red represents significant bins unique to the component, blue those common to both. **D:** As C but for PI component. **E:** TF motif enrichment (foreground versus background) around expressed CAGE-derived promoters associated with GM12878 or HeLa-S3 biased differentially expressed PD or PI bins. See also Supplementary Figure S2E.

To examine the localised patterns observed for expression levels not attributable to position (PI component, Fig. 2A), we trained a random forest model (Methods) on GM12878 transcriptional components to predict the presence or absence of DNase I hypersensitive sites (DHSs), histone variant H2A.Z and post-translational histone modifications H3K4me3 and H3K27ac (binarised DNase-seq and ChIP-seq data (Zerbino et al., 2015) in each bin), each associated with features of (transcriptionally) active promoters (The ENCODE Project Consortium, 2012). The resulting models allowed us to generate a probability distribution for each mark given each transcriptional component. For all tested marks we observed a clear bias with stronger predictive power from the PI component than the PD component (Fig. 3B). These results indicate that the PI component, in contrast to the PD component, carries information about promoter-localised and expression-level associated effects.

Overall, we found that both the PD and PI components explained considerable proportions of expression levels in GM12878 cells (Supplementary Fig. S2A). Each component explained on median roughly half of the expression levels in expressed bins, with contributions varying between loci (Supplementary Fig. S2B). When compared with HeLa-S3 cells, we observed clear differences in both the PD and PI components between cell types and that differences were orthogonal between components (Fig. 3C,D). Differentially expressed bins in the PD component (Supplementary Table S2) were, in line with its relationship with chromatin compartments, to a large degree associated with cell-type differences in chromatin states, changing from silent to H3K36me3-associated active chromatin in up-regulated bins (Supplementary Fig. S2C-D). The PI component, on the other hand, showed localised differential expression of bins (Supplementary Table S3) that were associated with cell-type specific enrichments of predicted transcription factor (TF) binding sites from sequence motifs (Fig. 3E and Supplementary Fig. S2E), for instance NFKB and IRF in GM12878 cells.

Similar patterns of transcriptional components derived from GM12878 CAGE data were found when we applied transcriptional decomposition, guided by CAGE-estimated hyperparameters, to GM12878 RNA-seq data (The ENCODE Project Consortium, 2012) (Supplementary Fig. S3, Supplementary Table S1, Methods), confirming the observed characteristics and differences between CAGE-derived components. Taken together, we conclude that expression data can be decomposed into a positionally dependent component, revealing chromatin compartment activity, and a positionally independent component carrying information about localised, independent expression-associated events.

### Expression associated domains show clear resemblance with active chromatin topology

Apart from displaying clear shifts at compartment boundaries, we noted that the PD component contained sub-patterns within broader consecutive regions of positive signals (Fig. 2A). We posited that such structures could represent finer, transcriptionally active chromatin organisation not necessarily reflecting chromatin compartments, but rather boundaries of active TADs. To test this hypothesis, we trained a generalised linear model (GLM) to predict HiC-derived TAD-boundaries (Rao et al., 2014) from features derived from transcriptional components (Supplementary Table S4, Methods). The GLM yielded an area under the ROC curve (AUC) of 0.73 (AUC 0.85 in regions of positive PD signal), indicating that there is information in expression data to infer TAD boundaries (Supplementary Fig. S4A,B). Among features considered, the model ranked the PD gradient (first order derivative), the PD inter-cell stability, and PD variance among the most important for predicting TAD boundaries (Supplementary Fig. S4C). Based on the GLM feature importance ranking, we devised a score to rank PD boundaries at significant gradients in the PD signal that also had low positional standard deviation (high stability) across cells (Fig. 4A, Methods). In GM12878 cells, we detected 4,158 boundary locations of PD sub-patterns. We refer to the regions demarcated by PD boundaries as expression associated domains (XADs, Fig. 4A).

**Figure 4.**
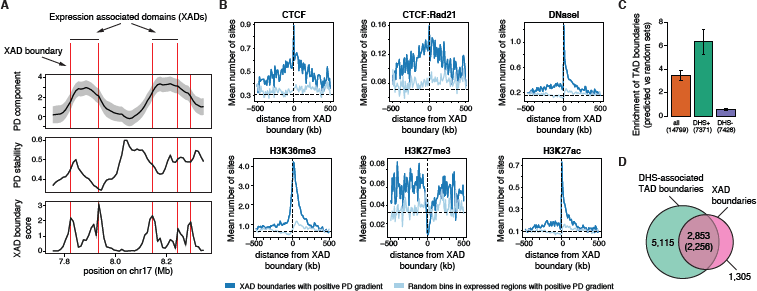
Expression associated domains mark regions of active topological domains. **A**: Approach for identifying boundaries of expression associated domains (XADs) based on a PD boundary score. Shown are PD signal (mean +/- standard deviations), PD stability (across cell PD standard deviation) and the XAD boundary score. **B**: Average GM12878 profiles of binarised ChIP-seq data for CTCF, CTCF in combination with Rad21 (cohesin), DNaseI, H3K36me3 H3K27me3, and H3K27ac at XAD boundaries with positive PD gradient (dark blue) and at random expressed bins with postive PD gradient. Dotted vertical line represents boundary locations and horizontal dotted line represents background mean for given ChIP-seq mark. **C**: Enrichment of GM12878 TAD boundaries among XAD boundaries compared to random bins proximal to expressed bins (DHS+ for DHS associated, DHS- for DHS non-associated). **D**: Venn diagram of association between GM12878 XAD boundaries and proximal (within 5 bins) DHS-assocated TAD boundaries.

We next assessed the occurrence of DHSs, ChIP-seq binding site peaks for architectural proteins CTCF and Rad21 (a subunit of cohesin) and histone modifications H3K36me3, H3K27me3, and H3K27ac around GM12878 XAD boundaries (Fig. 4B). We observed an enrichment of DHSs, H3K27ac, and binding of CTCF alone and, albeit weaker, in combination with Rad21 around XAD boundaries. In addition, H3K36me3, H3K27ac, and DHSs were more enriched downstream than upstream of positive-gradient XAD boundaries. The opposite trend was observed for H3K27me3. The observed DHS and ChIP-seq patterns around XAD boundaries (Fig. 4B) resemble those around TAD boundaries in active compartments (de Wit et al., 2015; Li et al., 2013; Pope et al., 2014; Tang et al., 2015). In line with these and the GLM results, we observed that GM12878 XAD boundaries were in general highly enriched in HiC-derived TAD boundaries (Rao et al., 2014) from the same cells (>3-fold enrichment over background, Fig. 4C). Specifically, XAD boundaries were highly enriched in DHS-associated TAD boundaries (>6-fold enrichment over background), but not in those distal to DHSs (Fig. 4C and Supplementary Fig. S5A,B). Out of 4,158 GM12878 XAD boundaries, 2,853 (69%) were proximal (within 5 bins) to DHS-positive TAD boundaries (Fig. 4D). Still, 69% (5,115 out of 7,371) of DHS-positive TAD boundaries were distal to XAD boundaries, suggesting that XADs represent a subset of DHS-positive TADs. Joint XAD and TAD boundaries were primarily found within active chromatin while XAD-unsupported DHS-positive TAD boundaries more frequently resided in facultative or constituitive heterochromatin (Supplementary Fig. S5C, p<1e-258, Chi-squared test). Furthermore, joint XAD and TAD boundaries displayed a higher HiC (Rao et al., 2014) chromatin interaction directionality (Dixon et al., 2012) than DHS-positive TAD boundaries distal to XAD boundaries (Supplementary Fig. S5D,E), indicating that expression-associated chromatin is linked with stronger TAD boundaries (greater insulation). However, both sets were similarly associated with Rad21 (Supplementary Fig. S5F,G), whose co-binding with CTCF is believed to provide strong TAD boundaries (de Wit et al., 2015; Li et al., 2013; Tang et al., 2015). Taken together, these results demonstrate that expression data (PD component) can be used to infer chromatin topology in active chromatin compartments.

### Transcriptional components are informative of regulatory chromatin interactions

Since positionally attributable RNA expression was strongly associated with structures of transcriptionally active TADs, we questioned the utility of expression data alone in reflecting individual proximity based interactions. Namely, what does it mean to be proximal in the nucleus, from a transcriptional perspective?

To this end we collected a total of 25 features in GM12878 that may be derived from expression datasets alone (Supplementary Table S5), relating to the values, differences, cross-cell type correlations, standard deviations and stabilities of the decomposed components, as well as those relating to CAGE-measured promoter and enhancer activities, directionality scores, XAD boundary insulation and chromosomal distance. To test the power of expression-universal features to classify chromatin interactions, we used a random forest classification scheme where we compared to GM12878 promoter-capture HiC (CHiC) data (Mifsud et al., 2015) at 10kb resolution (Methods). We used a lenient interaction score threshold to define positively interacting bin-bin pairs (bin-bin distance > 50kb and ≤ 2Mb, based on CHiCAGO (Cairns et al., 2016) score ≥ 3, see Table S6), for which each pair referred to a CHiC promoter bait and a potential target that overlapped a transcribed promoter (FANTOM Consortium and the RIKEN PMI and CLST (DGT) et al., 2014) or transcribed enhancer (Andersson et al., 2014). In order to deal with the resulting unbalanced data, we over-sampled (Chawla et al., 2002) the positives and under-sampled the negatives in the training data to a fully balanced set across distances in a ten-fold cross-validation scheme. In each cross-validation round, we balanced the training data and predicted on held-out data at 20:1 or raw (unmodified) negative:positive (NP) ratios.

Overall, we observed a good predictive performance (AUC: 0.74) on raw NP ratios for lenient interaction thresholds (Fig. 5A). Using a strict interaction threshold (CHiCAGO score ≥ 5) for evaluation increased the AUC (0.83) but reduced the recall (Fig. 5B). At a 20:1 NP test ratio, the overall predictive performances increased (AUC 0.82 and 0.88 for lenient and strict interaction thresholds, respectively). Interestingly, we observed a better precision and recall when evaluation of results was limited to enhancer targets (Supplementary Table S7, AUC of 0.85 and 0.89 for lenient and strict interaction thresholds at a 20:1 NP ratio, respectively; see Supplementary Fig. S6 for the effect of interaction thresholds on EP prediction performance). We observed varying predictive performances across chromosomes (Supplementary Fig. S7), likely reflecting differences in gene densities and transcriptional activities (data not shown). Despite the overall good EP predictive performance, both precision and recall decreased over increasing distances on held-out test data, in accordance with an increasing NP ratio (Supplementary Fig. S8). To circumvent the distance effect, we established a distance-dependent threshold in random forest voting, guided by the optimal F1 score, significantly improving the predictive performance over distance (Supplementary Fig. S9). Taken together, we conclude that there is a wealth of information from properties of expression data alone which could explain chromatin interactions, and in particular enhancer-promoter interactions. This suggests a tight coupling between expression of regulatory elements and their proximity.

**Figure 5.**
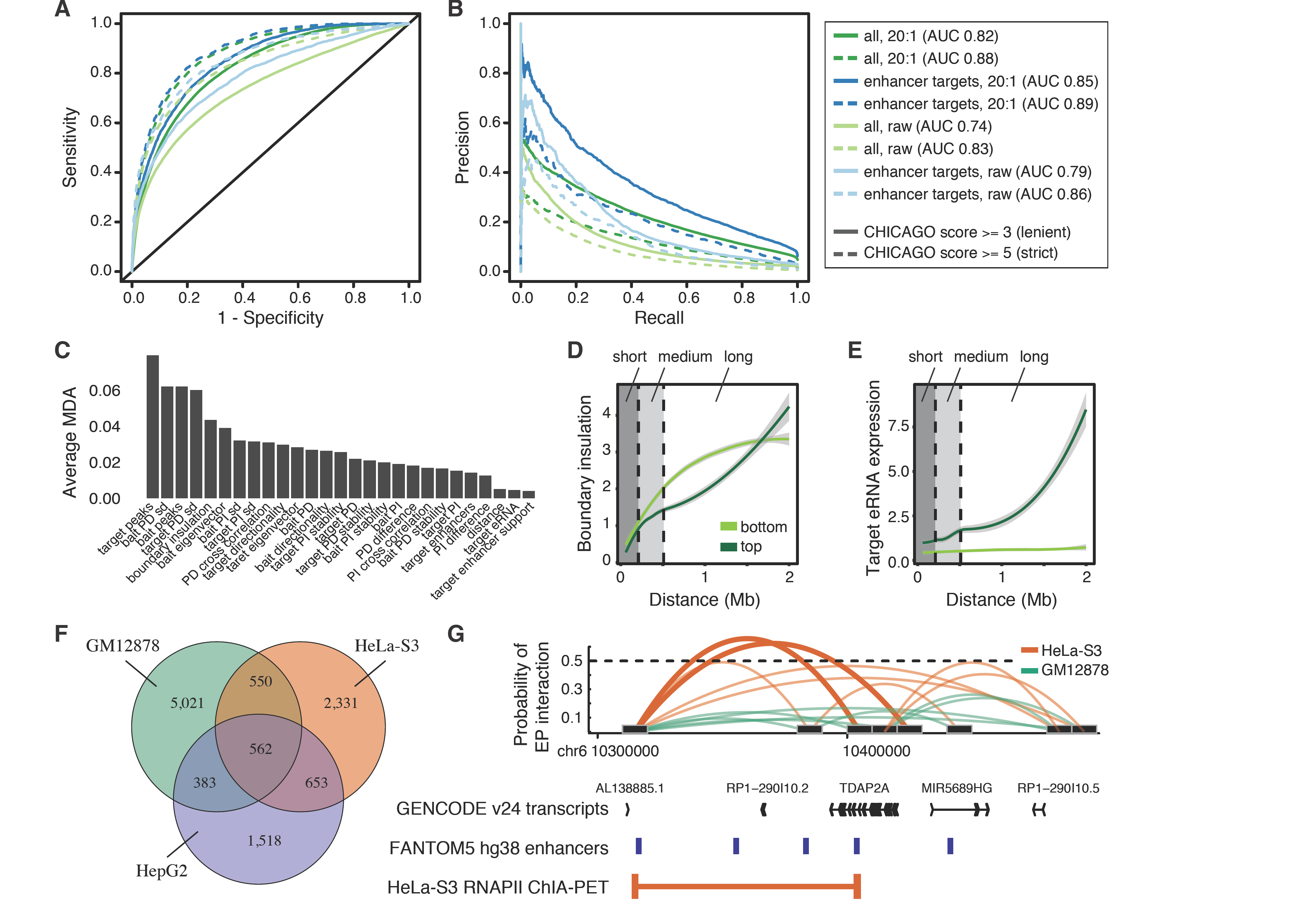
Expression data is predictive of cell type specific regulatory interactions. **A-B:** Performance curves for predicting bait-target interactions from CAGE-universal features, split according to CHICAGO score, testing negative to positive ratios and target feature type. **C:** Features predictive of bait-target interactions, ordered by average mean decrease accuracy (MDA) across models from 10-fold cross validation. **D,E:** Loess curves representing feature separation over distance between high (top) and low (bottom) predicted probabilities, shown for features (**D**) XAD boundary insulation *(nbounds)* and (**E**) enhancer expression at target *(eRNA_targ).* **F**: Overlaps of predicted bait-enhancer interactions between GM12878, HeLa-S3, and HepG2 cells. **G**: An example of a loop predicted in HeLa-S3, but not in GM12878 cells, validated by HeLa-S3 RNAPII ChIA-PET interaction data.

Next we asked what properties of expression data explained chromatin interactions. High transcriptional activity at the target or bait was ranked among the top features for predicting chromatin interactions (Fig. 5C). In addition, the standard deviation of the PD component at either the bait or the target was informative (Fig. 5C). Predicted interactions had a lower PD standard deviation on average (Supplementary Fig. S10), suggesting that higher confidence interactions are associated with more stable expression levels. Boundary insulation, defined as the number of XAD boundaries predicted between the bait and the target, also ranked highly, with a smaller insulation associated with a higher probability of a chromatin interaction compared to pairs at a similar distance (Supplementary Fig. S10). Since the observation of this property forms the definition of TAD domains, this lends support of XAD boundaries as predictors of TAD boundaries. Separately training three random forest classifiers for short, medium, and long-range distances (covering bait-target distances within (50,200], (200,500], and (500,2000] kb, respectively) did not improve the predictive performance compared to the full model (data not shown), but allowed us to further investigate features driving long-range interactions (Supplementary Fig. S11A). As expected, boundary insulation had a higher influence on long-distance interactions than shorter ones (Fig. 5D). Interestingly, when we specifically considered interactions between enhancers and promoters, eRNA expression at the target enhancer clearly distinguished predicted positive from predicted negative interactions, and its importance increased over increasing distances (Fig. 5E).

Motivated by the good performance in predicting enhancer-promoter (EP) interactions, we next used the GM12878-trained random forest model to predict EP interactions also in HeLa-S3 and HepG2 cells (Supplementary Tables S8, S9). We noted that many predicted interactions were specific to each cell type (Fig. 5F, 48-78%), with only a small fraction (9-20%) of interactions shared between all three cell lines (see Supplementary Fig. S12 for results using a strict interaction threshold). For instance, the promoter of gene *TDAP2A* was predicted to interact with a ~100kb downstream enhancer specifically in HeLa-S3 cells, supported by HeLa-S3 RNAPII ChIA-PET interaction data (The ENCODE Project Consortium, 2012) (Fig. 5G).

In support of eRNA production at positive interactions, both enhancers and promoters in predicted cell-type specific interactions clearly showed an expression bias towards the cell type in which the interactions were identified, in contrast to shared EP interactions (Supplementary Fig. S11B,C). These results indicate that expression of regulatory elements is both reflecting their cell-type specific regulatory activity and their regulatory interactions. This is supported by observations that regulatory active enhancers which are interacting with promoters are more likely to be transcribed than non-interacting ones (Sanyal et al., 2012). Taken together, we demonstrate that transcription is highly informative of three-dimensional chromatin architecture and may be used to accurately infer EP interactions.

### Transcriptional decomposition reveals regulatory differences between human cell types and guides interpretation of disease-associated genetic variants

We have above demonstrated that positionally attributable expression (PD component) can be used to accurately infer boundaries of active TADs as well as differences in chromatin compartments and EP interactions between well-established and biologically distal cells lines. We continued with exploring what insights could be gained by transcriptional decomposition of primary cell CAGE data (FANTOM Consortium and the RIKEN PMI and CLST (DGT) et al., 2014), for cell types for which chromatin topology is to a large degree unknown. We applied transcriptional decomposition to replicated primary cell CAGE data (FANTOM Consortium and the RIKEN PMI and CLST (DGT) et al., 2014) expanding to a total of 76 human decomposed cell types from 249 CAGE libraries (Supplementary Table S11, Methods). Using the resulting components, we calculated XAD boundaries and extended the previous set of pairs defined in the GM12878 training above to a common set of bin-pairs across all cell types (Supplementary Table S10), over which EP interactions were predicted.

Consistent with observations in cell lines (Fig. 2A), the PD and PI components displayed opposing trends when compared across cell types. Many genomic bins showed highly similar PD signal across cells, while the PI component was rarely shared across more than a few cell types (Supplementary Fig. S13A,B). The PI component grouped more closely according to cell type association than the PD component and clearly distinguished blood cells from mesenchymal, muscle, and epithelial cells (Fig. 6A,B). XAD boundaries tended to be either highly cell-type specific or ubiquitous (Supplementary Fig. S13C), similar to that of TAD boundaries (Schmitt et al., 2016), reflecting the cell-type specific behaviour of features around XAD boundaries (Fig. 4B), e.g. DHSs (Thurman et al., 2012). In comparison, EP interactions exhibited very little agreement across cell types (Fig. 6C), with very few shared interactions across the full repertoire of samples (Supplementary Fig. S13D). Interestingly, cell type specific EP interactions were on average more distal than interactions shared across cells (Supplementary Fig. S13E). When examining EP interactions between groups of cells we observed clear differences at key identity genes. Leukocyte biased genes *ARHGAP25* (Fig. 6D) and *CD48* (Supplementary Fig. S14) showed clear differences in predicted EP interactions, which were validated by GM12878 CHiC interaction data. Taken together, these results indicate largely invariant chromatin compartments between cells and that cell-type identities are governed by differences in EP interactions or position-independent effects, which may be driven by localised, cell-type specific TF binding events.

**Figure 6.**
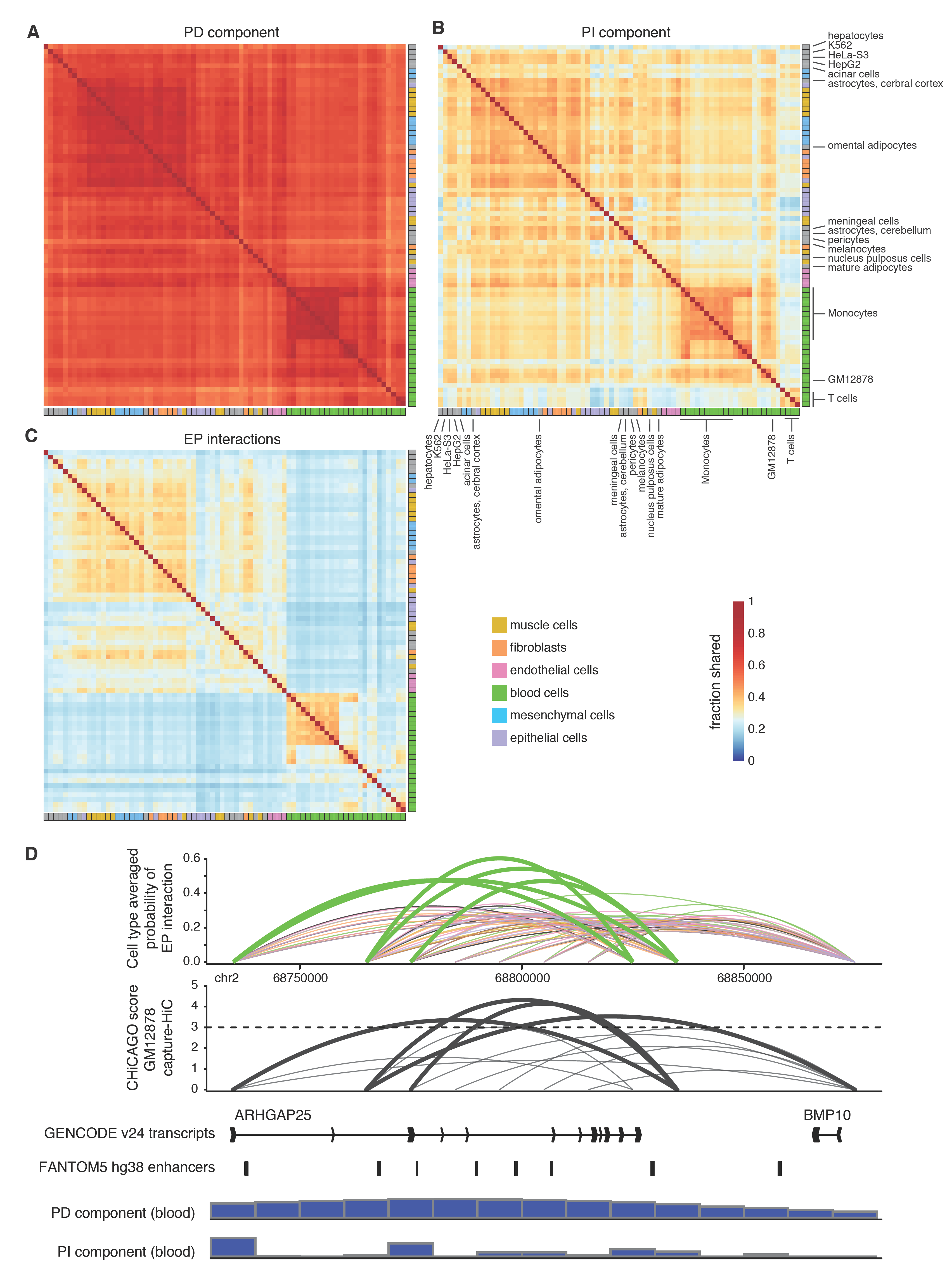
Transcriptional decomposition across 76 human cell types. **A-C**: Heat maps depicting pairwise similarities of the PD (**A**), PI (**B**) components, and EP interactions (**C**) across cell types. All similarity scores calculated between cell types using 1 — *L*_1_-norm on binary data sets based on the sign of the PD, PI components or the presence/absence of EP interactions. **D**: Predicted probability of EP interactions averaged across groups of cells and CHiCAGO score of interaction based on GM12878 capture HiC data. Below are displayed GENCODE v24 transcripts, FANTOM5 enhancers, the average PD and PI component across blood cells.

Many complex genetic diseases are associated with genetic variants outside of protein-coding gene sequences in gene-distal enhancers (Andersson et al., 2014; Maurano et al., 2012) and these variants have been suggested to explain a large portion of disease heredity, in particular in immunological disorders (Finucane et al., 2015). Thus, we investigated the utility of decomposed expression data and predicted EP interactions in the interpretation of disease associated genetic variants. Based upon enrichment analysis of trait-associated SNPs (Welter et al., 2014) and those in strong linkage disequilibrium (Methods), we observed largely variable preferences between traits and transcriptional components (Supplementary Fig. S15A). Interestingly, several diseases and traits showed a preference for enrichment in positionally attributable expression (PD positive bins) or within 2 bins of XAD boundaries, indicating that associated genetic variants may alter chromatin compartments or the activities or structures of enclosing TADs. Furthermore, whilst most trait enrichments were restricted to a few cell types, many of those biased to positionally attributable expression, for instance Crohn’s disease, displayed broad association across the panel of investigated cell types (Supplementary Fig. S15B). Monocytes, including those subjected to various pathogens, T cells, B cells and natural killer cells were found amongst the most highly ranked cell types associated with Crohn’s disease in PD and PI components as well as at XAD boundaries (Fig. 7A). This is consistent with the strong link between Crohn’s disease, inflammation, innate and adaptive immune system deficiency (Torres et al., 2017). Amongst genes in SNP-associated bins, we found Crohn’s disease-associated genes *STAT3, ATG16L1,* and MHC genes *HLA-DWB, HLA-DRA,* and *HLA-DQA2* (McGovern et al., 2015). Similar to Crohn’s disease, lymphoid leukemia was strongly associated with cells of the immune system (Supplementary Fig. S16A), but had different genes associated with the PD and PI components and at XAD boundaries, including *IRF4, IKZF1*, *ARID5B, CEBPE, IRF8,* and *GCSAML.*

For Crohn’s disease, we found a significant enrichment of enhancer-overlapping SNPs in predicted EP interactions of monocytes treated with Cryptococcus and Salmonella, as well as in peripheral blood mononuclear cells (Fig. 7A). SNP-associated enhancers were predicted to interact with several disease-relevant genes, including *PTGER4* (Fig. 7B), *TRIB1* (Fig. 7C), *CCL1, OTUD1, ARMC3,* (Franke et al., 2008; McGovern et al., 2015), and genes so far not associated with Crohn’s disease *ENKUR,* and *THNSL1*. For example, promoters of *PTGER4* were predicted to interact with enhancers more than 250kb upstream of the gene situated in an LD block of GWAS-associated SNPs (Fig. 7B). Similarly, promoters of *TRIB1* were predicted to interact with a ~100kb downstream enhancer overlapping a disease risk locus of SNPs (Fig. 7C). These results clearly demonstrate how transcriptional decomposition of expression data can be used to gain insights into disease etiology, and thus the value of our generated resource.

**Figure 7.**
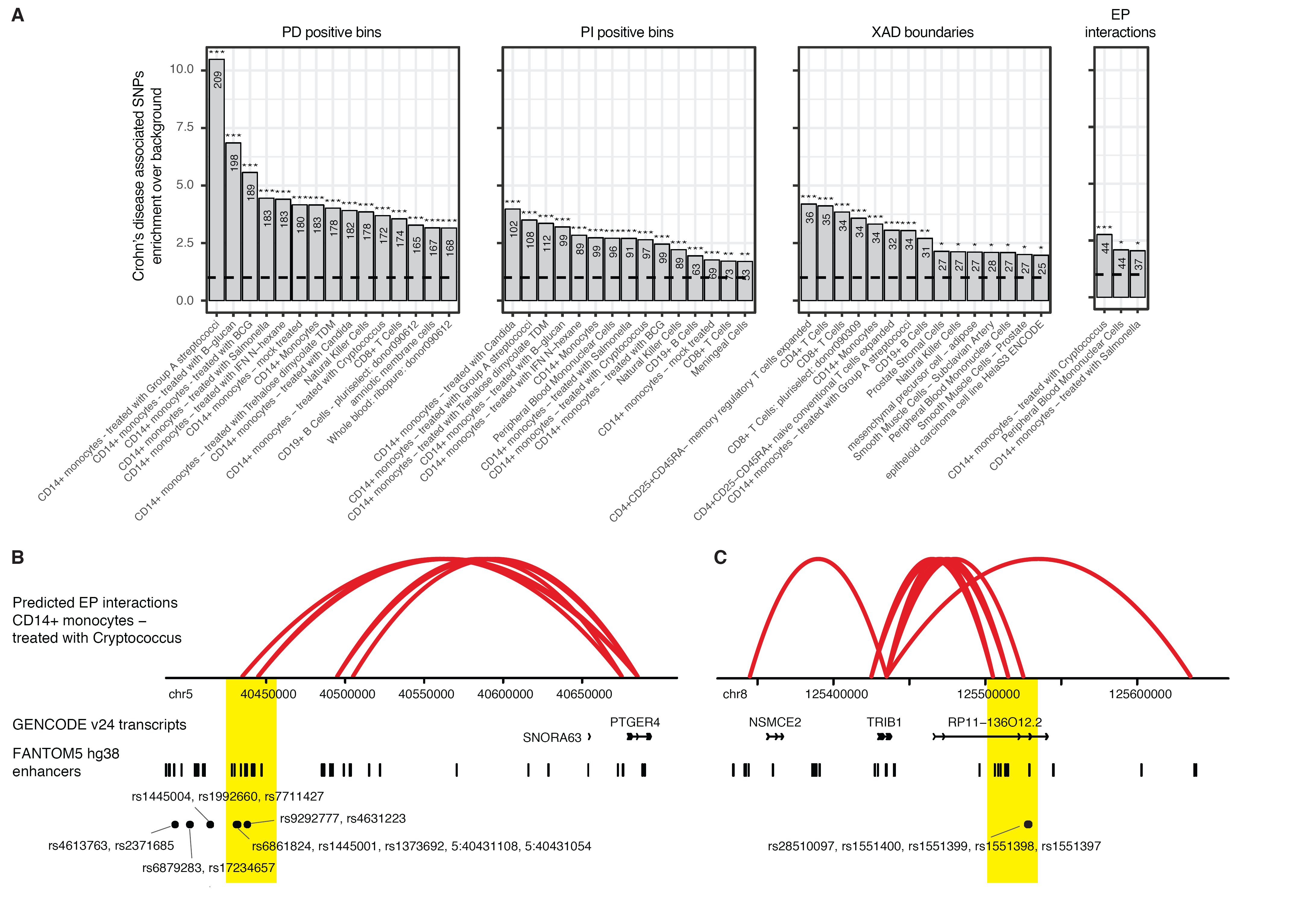
Analysis of Crohn’s disease associated SNPs reveals cell-type preferences and regulatory associations. **A**: Crohn’s disease SNP enrichments in transcriptional components, XAD boundaries, and inferred enhancer-promoter interactions (FDR-corrected *χ*^2^ tests based on the PD/PI positive bins or presence of XAD boundaries/target enhancers per cell type, and trait associated SNPs). See also Supplementary Figure S16 for enrichments in lymphoid leukemia. **B,C**: Predicted EP interactions in CD14+ monocytes treated with Cryptococcus, for genes *PTGER4* (**B**) and *TRIB1* (**C**) for which interacting enhancers overlap or are in close proximity with disease-associated SNPs (highlighted in yellow). Displayed are (from top to bottom) predicted EP interactions, GENCODE transcripts, FANTOM5 enhancers, and the locations of relevant SNPs.

## Discussion

A detailed understanding of the intricate relationship between transcription and three-dimensional chromatin organisation and to what extent they are attributable to each other has been lacking. Under the assumption that the link between transcriptional activity and chromatin organisation is reflected, at least partly, by chromosomal positioning, we have in this study aimed to separate the fraction of RNA expression that is associated with chromatin topology from localised, independent effects. To this end, we have developed a transcriptional decomposition approach that decomposes expression data into positionally dependent and independent components along chromosomes. We show that positionally attributable expression, according to the PD component, closely reflects chromatin compartments and domain architecture, and accounts for a sizeable overall fraction of TU expression levels. This points to the existence of large constraints on genomic organisation, suggesting that the global maintenance of chromatin formation is crucial for correct and precise transcriptional output. On the other hand, positionally independent expression, according to the PI component, carries information about promoter-localised and expression level associated effects. We suggest that positionally independent effects may be the result of cell-type specific programmes involving for instance the differential binding of transcription factors, potentially based on a layer of localised regulation not necessarily attributable to higher order three dimensional chromatin. However, we observe that the level of positional dependency of expression data varies between loci and cell types, suggesting locus- and cell-dependent effects of topological organisation and activity as well as varying impact of individual gene regulatory programs. This suggests that genes strongly associated with non-positional expression may be more resistant to perturbations in chromatin domains, such as deletions of TAD boundaries, particularly in the context of a certain disease.

The usage of expression data to infer topological chromatin organisation is limited by the inability to inform on transcriptionally silenced, closed or poised states. However, by focussing only on expressed TUs, such as those observed with CAGE, and their relative relationships, we can attempt to understand patterns that are highly relevant to various cell types of interest. For example, the majority of detected XAD boundaries were proximal to TAD boundaries associated with an open chromatin site in GM12878 cells. Interestingly, XAD boundary locations were seen to be strongly informed by the presence of stable PD signal across cell types, reflected in their high degree of sharing, which closely corresponds to the nature of TADs, whose locations appear largely cell type invariant (Dixon et al., 2012; Schmitt et al., 2016). This is likely connected to the enrichment of TSSs for actively transcribed housekeeping genes at the locations of TAD boundaries (Dixon et al., 2012; Hug et al., 2017), whose expression levels remain stable across cell types. Our predictions also suggest that active transcription at open chromatin loci is linked with stronger TAD boundaries, according to the observation that chromatin interaction directionality scores were stronger at TAD boundaries proximal to XAD boundaries compared to those distal from XAD boundaries, which were associated with closed chromatin. These results suggest that the presence of active transcription aids in the reinforcement of the insulation properties associated with boundary formation, but may also point to different regulatory mechanisms or roles involved in boundaries according to their link to active transcription.

We have shown that expression data alone is predictive of fine-scale chromatin interactions, based on predictive models incorporating expression component information, expression strength, stability and distance. Our results concur with, and significantly extend, the previous finding that CAGE is an informative predictor of interactions (Andersson et al., 2014; Whalen et al., 2016). Crucially, we bring up the question of what proximity within the nucleus means from the stand-point of transcriptional initiation, and suggest the two to be tightly coupled. Analysis of predicted features point to a model whereby a strongly expressed TU with low expression noise within its positionally dependent component are key predictors of within-domain interactions in a given cell type, and whereby strength of target expression becomes more important at long distances. Interestingly, of all predicted chromatin interactions, those restricted to EP-interactions showed the strongest predictive power, were biased towards the cell types in which they are actively transcribed and target enhancer RNA expression levels rapidly increased with the distance between the bait and the target.

Since transcriptional decomposition is broadly scalable across large numbers of samples and applicable to different expression sequencing assays, including CAGE and RNA-seq, it potentially paves the way for large-scale cost- and time-effective computational analyses across atlases of high quality expression data sets, such as FANTOM5 (FANTOM Consortium and the RIKEN PMI and CLST (DGT) et al., 2014) or GTEx (GTEx Consortium, 2013). We have applied our models to 76 cell types from the FANTOM5 consortium, generating predicted components, XAD boundaries and EP-interactions. This resource allows a deeper understanding of the dynamic regulation at key identity genes in a wide diversity of cellular states not yet subjected to methods informing on their higher order structures. Comparison across cell types revealed extensive sharing of positionally attributable expression, while positionally independent effects and, in particular EP interactions, were to a large degree cell type specific, particularly at long distances, and separated cells into groups of similar type. These results indicate that chromatin compartments are mostly invariant across cells and that fine-tuning of cell-type specific expression levels is mainly mediated by promoter-localised, position-independent effects or enhancer regulation.

While many complex diseases have clear cell-type specific effects and positionally attributable expression were chiefly shared between cells, we found an unexpected preference for enrichments of trait-associated genetic variants in the PD component. This suggests that the etiology of certain diseases may be coupled with the disruption or alteration of TAD structures or chromatin compartments, which finds support in the literature (Franke et al., 2016; Krijger and de Laat, 2016). Other traits were associated with cell-type specific enrichments of variants in the PI component or at enhancers in cell-type specific EP interactions, likely to cause altered gene expression levels independent of chromatin structures. Our generated resource of regulatory interactions enabled us to identify several cases of predicted EP interactions for which enhancer overlap with diseases-associated SNPs were linked with promoters of genes often associated with the disease itself. Overall, we expect that our transcriptional decomposition approach and resource will have large implications for future interpretations of genetic variants associated with disease in cell types that are otherwise largely intractable.

## Acknowledgements

This project has received funding from the Danish Council for Independent Research and the European Research Council (ERC) under the European Union’s Horizon 2020 research and innovation programme (grant agreement no. 638173). The authors would like to thank Hideya Kawaji for remapping and DPI tag clustering of FANTOM CAGE data and members of the Robin Andersson lab and Albin Sandelin lab for rewarding comments and discussions.

## Author contributions statement

S.R and R.A. conceived the project; S.R conducted most analyses, with support from M.D, L.D and R.A; S.R and R.A wrote the paper; all authors reviewed the final manuscript.

## Additional information

Data generated in this project have been made available at https://zenodo.org/record/556727 (DOI:10.5281/zenodo.556727) and https://zenodo.org/record/556775 (DOI:10.5281/zenodo.556775). The authors declare no competing financial interests.

## Methods

Most analyses were carried out in *R* (R Core Team, 2016). All liftovers from hg19 to hg38 were carried out using the chain `hg19ToHg38.over.chain` downloaded from UCSC (Kent et al., 2002) using the R package `rtracklayer` (Lawrence et al., 2009), unless otherwise stated. All overlaps between genomic regions were carried out using the R package `GenomicRanges` (Lawrence et al., 2013), unless otherwise stated.

### Processing of CAGE data sets

#### Source of CAGE data

Data was produced by the FANTOM5 project (FANTOM Consortium and the RIKEN PMI and CLST (DGT) et al., 2014). The data was mapped to hg38 and CTSSs (CAGE tag starting site) were clustered into tag clusters (TCs) according to decomposition peak identification (dpi) generation as per FANTOM5 (Kawaji, 2017). Only samples with more than 0.5 million tags mapping within the TCs were included in the analysis.

#### Enhancer calling

Enhancers were called based on bidirectional balanced RNA signatures as per the FANTOM5 consortium (Andersson et al., 2014). Enhancers were only identified distal to known exons (+/-100bp region from boundaries) and transcription start sites +/-300bp, defined by GENCODE v24 annotation. In total, 63,285 enhancers were identified across 1,829 libraries.

Due to varying noise levels across FANTOM libraries and the intrinsic low expression levels of transcribed enhancers, library-specific noise levels were estimated to define of robust set of enhancers in each sample. For each library, expression was quantified in randomly sampled genomic regions distal to assembly gaps, DNase hypersensitive sites (ENCODE), known exons and gene TSSs (GENCODE) to create a genomic background expression distribution. For each library, we called an enhancer active (used) if its expression was above the 99.9th quantile of the library’s genomic background expression distribution. The robust set of enhancers consist of 60,215 over 1,829 libraries, being significantly expressed in at least one library. The expression was quantified and TPM (tags per million) normalised according to the total number of mapped reads within the full set of TCs.

#### Data binning into 10kb regions

CTSS files containing positions of raw CAGE counts from libraries for the ENCODE tier 1 cell lines (GM12878, K562, HeLa-S3 and HepG2) were intersected with non-overlapping 10kb regions across the genome. For each chromosome (chr1-chr22 and chrX), regions were defined from coordinate 1 (1-based) and in consecutive complete 10kb blocks, up to two blocks after the last bin containing a single CAGE tag across the set of libraries. Region sizes of 40kb and 100kb were also considered, but 10kb was exclusively used in the analyses for comparability to high resolution chromatin capture data sets.

#### Data distribution

Due to the sparsity of transcription in bins associated with non-structural regions and regions poorly mapped, including the case of libraries with poor sequencing depth, a zero-inflated negative binomial distribution was assumed to hold for the counts across the 10kb bins on each chromosome, given by, for bin *i, Y_i_* ~ ZINB(*p_i_, μ_i_, k*), where *Y_i_* is bin count, *p_i_* represents the zero probability parameter, *μ_i_* represents the mean and *k* the size, or dispersion, parameter. The mean is given by 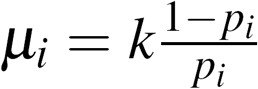, with variance 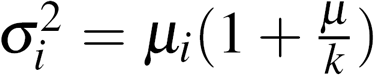. The model assumes a zero-truncated negative binomial distribution on the non-zero counts (*Y_i_ ~ NB*(*μ_i_, k*) for *y_i_* = 1,2,…) with probability equal to 1 – *p_i_*.

#### Decomposition using random effects model

The decomposition models the mean log count as a combination of an intercept and two random effects, PD and PI, set up as *v_i_* = *α* + PD_*i*_ + PI_*i*_ where *α* is the intercept, PD_*i*_ and PI_*i*_ are random effects for the PD and PI components respectively and *v_i_* is the linear predictor given by *v_i_* = log(*μ_i_*) – log(*E*) for library depth *E* and log(*E*) is the offset term. The PD component is modelled as a first order random walk 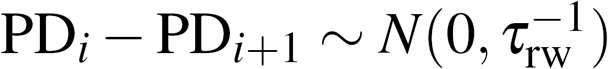 where PD_*i*_ – PD_*i*+1_ represents the component difference between successive bins and *τ*_rw_ is the precision of the normally distributed differences, with mean 0. The PI component assumes that bins are represented by a vector of independent and Gaussian distributed random variables with precision *τ*_*iid*_.

#### Model fitting

The model was fit in the form of a Bayesian mixed model using the software *R – INLA* (Rue et al., 2009) which implicitly assumes a Gaussian field on the parameter space and uses a Laplacian approximation to allow for fast and deterministic convergence of parameters. Hyperparameters are defined for the size and zero probability parameters (gamma distributed and gaussian dstributed prior distributions respectively) of the negative binomial distribution, and precision parameters *τ*_*rw*_ and *τ*_*iid*_ (log-gamma distributed priors) for the PD and PI components respectively. Three replicates for each of the ENCODE cell lines were included, assuming library depths equal to the sum of the tags in their respective CTSS files. ENCODE cell lines were modelled con-currently for each chromosome, assuming a common distribution for the hyperparameters.

#### Model scaling and convergence

In order to allow for efficient prior estimations we scaled the random walk components such that the average variance (measured by the diagonal of the generalized inverse) is equal to 1. In order to achieve efficient convergence and avoid precisely defining priors beforehand, the option `diagonal=1` was set within the INLA call (to avoid falling into sparse errors). The posteriors based on the converged model were then fed into a second model specifying `diagonal=0` in order to achieve more accurate estimates.

#### Applying the model to RNA-seq data

To see if chromosomal transcriptional decomposition is broadly applicable to RNA datasets, we applied the same modelling procedure to deeply sequenced GM12878 RNA-seq samples (https://www.encodeproject.org/experiments/ENCSR843RJV/) (The ENCODE Project Consortium, 2012). The libraries were mapped to hg38 using hisat (Kim et al., 2015), using default parameters, multimappers were removed using samtools (Li et al., 2009) and reads were binned at 10kb resolution (based on bins defined previously for CAGE) using bamCoverage (Ramírez et al., 2014). Transcriptional decomposition was applied to the resulting counts, assuming library depth to be the total of the genome-wide bin counts. A range of hyper-parameters for the PD and PI components was tested (corresponding to the CAGE defined values of the parameters, and -3 to +3 relative) and the combination selected, per chromosome, whereby the PD component correlated the most strongly with the PD component in GM12878 CAGE (ignoring regions not within 25 bins of a transcription unit in RNA-seq).

### Analysis of PD and PI datasets

#### Proportions of transcription units represented by PD and PI components

To address the proportions of total mRNA levels allocated to each of the PD and PI components, the raw CAGE counts in GM12878 were used to identify 10kb bin regions with a mean count greater than 50 tags across the three replicates, thus ensuring most selected bins were positive in both components (but removing those which were not). For each of these bins the fraction 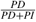 was calculated, where *PD* and *PI* are the respective estimates of the PD and PI components in the bin. These fractions were plotted as a histogram and the median value was identified as an average ballpark for relative allocation to PD component vs the PI component.

#### ChIP-seq data for ENCODE cell lines

ChIP-seq GRCh38 bigwig and peak data were downloaded from Ensembl FTP and Ensembl biomart based on Ensembl regulatory evidence v 84 (Zerbino et al., 2015).

#### Compartment data

Compartment coordinates for GM12878 (Rao et al., 2014) were lifted over from hg19 to hg38 using liftOver tool with default settings. For simplicity of interpretation, the five compartments types from (Rao et al, 2014) (Rao et al., 2014) were associated with Active (A1,A2), Facultative (B1) and Constitutive (B2,B3) chromatin environments.

#### Association between PD and chromatin environment

Overlaps between the 10kb bin regions and compartment regions were used to assign bins to chromatin environments, removing bins not having a corresponding overlap. Bins representing shifts between two different chromatin environments were deemed boundary bins and for sets corresponding to each possible shift (e.g. active to constitutive), the mean of the PD component signal was calculated for bins at an increasing distance either side of the boundary bins, in steps of 10kb up to and including 500KB. Cases at a given distance whereby another boundary bin was encountered were removed. For each set, a random sampling was used to generate a list of random boundary bins of length equal to the set size. Background sets were each generated 100 times to form distributions.

#### Correlations and differences within and between compartments

Intra-chromosomal bin pairs which overlapped with an annotated compartment were assigned an integer according to how many boundaries compartment boundary bins (see above) were between them (boundary insulation). For all bin pairs, both the absolute first order difference in the PD signal and the correlation between the four ENCODE cell lines was calculated. The differences and the correlations were averaged using the median either at each distance from 10kb apart to 2MB apart or across all distances, separately for each possible boundary insulation.

#### ChIP-seq biases in PD vs PI components

For each ChIP-seq binding mark (DNase1, H2AZ, H3K4me3, H3K27ac), 10kb bin regions were overlapped with binding locations to give the presence or absence of the mark in each bin. For each of the PI and PD components, the bin estimate, first order difference between bin estimates, standard deviation of the bin estimate, stability of the bin estimate (standard deviation across cell type PD component standard deviations) were calculated. For the two components separately, a random forest model was trained with the ChIP-seq binding presence or absence as the response. Out of the bag probabilities from the models based on the PD data and the PI data were compared directly between the two components.

#### Differentially expressed (DE) bins between cell types in PD and PI components

Posterior estimates and standard deviations of the linear combinations representing the difference in the PD or PI component between GM12878 and HeLa-S3 equivalent bins were generated from the transcriptional decomposition models (described above). Since posterior distributions of the estimates are approximately Gaussian, approximate p-values from z-scores were generated in order to produce standardised scores for the differences. A Benjamini-Hochberg correction was applied according to the number of bins containing an active transcription unit (s.t. there were ≥ 10 tags across the three replicates in at least one of HeLa-S3 or GM12878), and using an FDR < 0.01 to generate a list of significant DE bins in each component.

#### Bias towards ChIP-seq binding in PD vs PI differentially expressed bins

H3K27me3 and H3K36me3 ENCODE Broad Institute bigwig data (from Ensembl Regulatory Build v 84) were quantified in 10kb genomic bins. The aggregated signal values in each bin were TPM normalised (accroding to all genomic bins). The TPM values for H3K27me3 and H3K36me3 were then inspected at DE bins between GM12878 and HeLa-S3 cells (see above).

#### Transcription factor targets in PD vs PI differentially expressed bins

Active regions within the DE PD and PI bins were tested for transcription factor binding enrichment using Homer (Heinz et al., 2010). Regions were defined as actively transcribed FANTOM5 DPI TCs (Kawaji, 2017), requiring > 1 tpm in at least two of the six libraries (three GM12878 and three HeLa-S3). DPI TCs were extended to regions of -500/+100 around TSSs. DU regions of the PI components with up-regulation in Hela-S3 or in GM12878 (HeLa-S3 PI, GM12878 PI), and DU regions of the PI components with up-regulation in Hela-S3 or in GM12878 (HeLa-S3 PD, GM12878 PD), were tested individually against the universe of actively transcribed TC regions. The tests were performed using a stranded search with GC bias correction. Significantly enriched known motifs (FDR < 0.01) were selected in each of the four tests and plotted together in a heatmap using the *R* package `pheatmap` (Kolde, 2015) with Euclidean clustering of transcription factors (columns).

### XAD boundaries predictions in ENCODE cell lines

#### List of TAD boundaries

TAD boundaries regions based on 1kb resolution HIC data in GM12878 were downloaded (Rao et al., 2014) and lifted over to hg38, requiring a 1:1 correspondence between regions defined in each of the two builds. Boundaries were assigned to 10kb bins based on overlaps. Adjacent bins with boundaries were dealt with by assigning the boundary to the bin with the largest overlap, so that no two adjacent bins contained boundaries, resulting in a total of 14799 distinct bin-sized regions containing TAD boundaries.

#### GLMnet model for finding features associated with TAD boundaries

Features generated per bin according to those listed in Supplementary Table S4. The set of 10kb bins containing HIC boundaries (see above) was extended either side by 1 bin to supply boundary regions to predict on. Due to lack of CAGE information in non-structural regions, the set of bins was reduced to regions with a potential for a boundary to be predicted by using only the set within 250KB of a bin containing 5 tags or more in GM12878 (replicate sum). The response (presence or absence of a TAD boundary with a bin) was generated in two ways, first for all bins within this set, and secondly under the requirement that the boundary had to sit in a positive random walk region.

In order to assess features which might distinguish bins containing or not containing boundaries, we fit a logistic regression model using the `glmnet` (Friedman et al., 2010) package in R. The predictors were scaled before modelling for generating scores of relative importance, using `caret` (Kuhn, 2016) package in R. To test the performance of the model at predicting boundary regions, a 2-fold cross validation was applied, where the data was randomly split into two equal sized parts and the total performance was assessed according to the combined predictions on the halves of the data which were held out of the modelling after the corresponding half had been trained on. ROC and precision-recall statistics were generated on a per-chromosomal basis, and for all of the chromosomes together, using the `ROCR` package (Sing et al., 2005) in *R* and plotted using custom functions.

#### Algorithm for detecting XAD boundaries from CAGE data

The top three features from the generalised linear model outlined in the previous section were selected, namely the PD component (*PD*), the difference in the PD component (*PD*_diff_) and the PD component stability (*PD*_stab_). Based on on these three features, the following algorithm was implemented for the detection of XAD boundaries:

1. Calculate 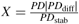.
2. Calculate local maxima of *X* and rank in order of largest to smallest values of *X*.
3. For each chromosome, calculate the proportion of bins of positive *PD, p_k_*, and take the top 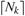 of the ranked maxima such that 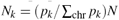, where *N* is the total target number of XAD boundaries.
4. Split the boundaries into “up jumps” and “down jumps” according to positive and negative values of *PD*_diff_, respectively.
5. Shift the “up jumps” right by one bin (to account for the discrepancy in which of the bins on either end of the *PD_diff_* should be called the bound. Leave the “down jumps” as is.
6. Return the vector sort(down jumps, up jumps).
7. Repeat above for the ENCODE cell lines.
8. For the final set of bounds in GM12878, choose the set such that the bound is in GM12878 + at least one other ENCODE cell line (and similarly for the set in the other cell types).

The above algorithm was applied to the ENCODE random walks, using the *EMD* (Kim and Oh, 2009) package in *R* to generate the local maxima and specifying a target of 5500 XAD. This resulted in a final list of 5109 XAD boundaries after the final filtering step.

#### Comparing XAD boundaries to ChIP-seq data

XAD boundaries were calculated at bins where the PD component has either a positive or negative change (first order difference in PD component). To generate enrichment of ChIP-seq marks around XAD boundaries, only those with a positive change are considered, in order to avoid biases from signal averaging. The results for the negative gradient boundaries are similar or opposing to that of the positive gradient boundaries.

For each of CTCF, DNaseI, H3K36me4, H3K27me3 and H3K27ac, the list of 10kb bins was overlapped with significantly bound sites to determine how many sites appeared in each bin. The mean number of sites overlapping the bins containing the predicted (positive change) XAD boundaries was calculated, then for bins one away from the boundary, and so forth in steps of 10kb up to a distance of 500KB. Cases at a given distance whereby another boundary bin was encountered were removed to avoid contaminated signal. For the background set, a set of random XAD boundaries of equal cardinality to the real boundaries were generated, under the conditions of positive gradient and within the set of bins plausible for being predicted as boundaries as previously defined. The same analysis was calculated for CTCF:Rad21, which was based on the number of CTCF sites multiplied by the number of Rad21 sites which overlapped a given bin.

#### Comparing boundary predictions to TAD boundaries

To calculate the overlap between the XAD set and the TAD set, regions around the XAD set were extended by 5 bins (50KB) on either side and then overlapped with the set of 10kb regions deemed to contain TAD boundaries. The number of TAD boundaries associated with a DHS site which fall within these regions were then counted, together with the number of predicted boundaries which fell in the vicinity of a DHS associated TAD boundary bin (defined as at least one DNase1 site from ChIP-seq overlapping the TAD boundary bin or one or more of the two adjacent bins).

To calculate the enrichment of XAD boundaries at the locations of boundary bins overlapping TAD boundaries, we calculated the proportion bins with predicted boundaries which overlapped the TAD boundary bin set and divided by the same value generated from a random set of boundaries (of the same length as the XAD set, falling within the set of plausible bins). We repeated the randomisation step 100 times and calculated the mean enrichment and standard deviation. The same analysis was repeated according to where the TAD boundary was DHS associated and where the TAD boundary was not associated with a DHS site (defined in the same way as with the overlaps above).

#### HiC directionality around XAD boundary predictions

Processed intrachromosomal HIC data for GM12878 at 10kb resolution (Rao et al., 2014) was downloaded and normalised, according to supplied recommendations, using the KR method. The directionality score (Dixon et al., 2012) was implemented and applied to locations of TAD boundaries whose corresponding liftovers in hg38 were either supported or not supported by XAD boundaries (within +/- 5 bins of the boundary). The Directionality score was calculated for +/-25 bins around the location of the boundary bins and for a jump size of 50 bins to calculate the direction bias of interactions. Means were plotted to obtain patterns of global directionality bias.

### Modelling of interactions in GM12878

#### Reprocessing of CaptureHIC data

CaptureHIC data for GM12878 was downloaded and fastq files extracted using the SRA toolkit (Leinonen et al., 2010). The data was mapped and corrected using HICUPS (Wingett et al., 2015), specifying bowtie2 (Langmead and Salzberg, 2012) for the mapping and GRCh38 for the genome (downloaded from ftp://ftp.ensembl.org/pub/release-85/). CHiCAGO (Cairns et al., 2016) was then applied to the remapped BAM files to call significant interactions against each bait, using default settings except for specifying 10kb binning and with the baits defined as the 10kb bin containing the defined sequences from the CaptureHIC protocol (Mifsud et al., 2015) lifted over from hg19 to hg38. Score cut-offs of ≥3 or ≥5 were applied to the final output in order to determine which bait-target pairs were considered as interacting, with the rest assigned as non-interacting.

#### Bait target pairs generation

Binned 10kb regions were overlapped with CaptureHIC protocol defined baits (Mifsud et al., 2015), lifted over to hg38. For each such ‘bait bin’, potential targets within 2MB (200 bins) were assigned and their numbers reduced according to the presence of CAGE tags in more than one replicate in at least one of the ENCODE cell lines. For analyses based on enhancer associated targets, targets were considered based on overlap with at least one CAGE defined enhancer which is active (more than one tag in more than one replicate) in at least one of the ENCODE cell lines.

#### Distance cut off

Only bait-target pairs which fall into the distance range of between 6 and 200 bins were considered (corresponding to greater than 50KB and up to 2MB).

#### Feature generation

The full list of features are listed in Table S5. Features from the PI and PD profiles were added for each bait-target pair separately for the bait and the target (see Table S4 for their descriptions). Enhancer information was added for the bait and targets based on the FANTOM5 enhancer set for hg38 (see above) - including the total eRNA output produced at the enhancer (replicate sum), the number of enhancers deemed active within the bin in the given cell line (at least one tag in at least one replicate) and the number of cell lines supporting the target enhancer (number of ENCODE cell lines with at least one enhancer active within the bin). Bin directionality was calculated based on pooled replicates, using a directionality score (Andersson et al., 2014) and assigning a value of 0 where no tags were present. The boundary insulation between bait and target was calculated according to the number of XAD boundaries observed in the intervening bins. Boundary insulation was defined up to 3; all values greater than 3 were given a score of 3. The peaks detected were the number of CAGE peaks overlapping the bait or target bin, according the hg38 DPI set from FANTOM5, which were transcribed in that cell type (at least one tag in at least one replicate). The cross correlation for the PI and PD components was calculated per chromosome by correlating the respective PI and PD profiles over four ENCODE cell lines between the bait and target bins, specifying Kendall for the correlation method. The first eigenvector was calculated from the cross correlation matrices for each chromosome and its value supplied for the bait and target bins. Distance was defined as the number of 10kb bins separating the bait and the target.

#### Dataset generation

Two datasets from GM12878 were analysed. For assessing the applicability of the model trained in GM12878 for predicting interactions in other cell types, the dataset as generated was kept in its current format (termed ‘raw’ format). In addition to this, since the number of positive interactions decays sharply with larger distances, a second dataset was generated where the ratio of negative to positive cases over distance was balanced by randomly sampling 20 negatives to each positive in the dataset, with replacement to account for cases where the negative rows were not at least 20 fold in number to the positive rows.

#### Data balancing for training sets

To generated a balanced dataset for training, SMOTE, as implemented in the *unbalanced* (Pozzolo et al., 2015) package in *R*, was applied to the data, specifying parameters *percOver=200* and *percUnder=150* to genere new positives together with under sampling the negatives to achieve a balance of 1:1 in the dataset. In order to balance the data set most fairly over distances, SMOTE was applied separately across each possible bait-target separation.

#### 10 fold cross validation procedure

The model was implemented in R using the `randomForest` (Liaw and Wiener, 2002) package sup-plemented with `foreach` (Revolution Analytics and Weston, 2015b) and `doParallel` (Revolution Analytics and Weston, 2015a) to run on multiple cores. We used 10 fold cross validation, whereby we split the dataset randomly into 10 equal sized pieces and held out a single piece as a testing set on each of the 10 runs of the model which was trained on the remaining 90%. All training was carried out based on a ChiCAGO score cut-off of 3. All performance statistics and probability estimates in GM12878 are based on predictions made across the held out runs over the full data set.

#### Splitting into short, medium and long models for feature importance

To assess whether there are different features which are important for specific distances without the bias of most examples being weighted towards short distances, the above analysis outline was repeated for the same data set but restricted to three possible distances: (50KB,250KB], (250KB,500KB] and (500KB,2MB]. For each set of distances, the MDA was averaged across the 10 runs in order to obtain a final feature performance.

#### Most efficient cut off for assessing predictions

To find the optimal probability cut-off for calling a predicted interaction, the value for which the F1 statistic was maximised was calculated using `optim` in *R*, according to the desired score cut-off. Since the most efficient cut-off is not fixed according to distance, the F1 statistic was maximised separately for five sets of bait-target distances: (50kb,100kb], (100kb,250kb], (250kb,500kb], (500kb,1MB], (1MB,2MB], and performance analysed for predictions generated using the resulting cut-offs. To calculate the effect of the CHiCAGO score cut-off on precision and recall, we optimised the F1 statistic separately for the five sets based on a range of score cut-offs (0.1, 0.25, 0.5, 1, 1.5, 2, 2.5, 3, 3.5, 4, 4.5, 5, 5.5, 6).

#### Assessing model performance

We used the pROC (Robin et al., 2011) package in *R* to generate AUC statistics and the caret (Kuhn, 2016) package in *R* to generate the precision, recall and F1 statistics. Plots were generated using custom functions based on statistics generated from the ROCR (Sing et al., 2005) package.

#### Generating predictions for HeLa-S3 and HepG2

The dataset for GM12878 was trained as described above against a score of ≥ 3 and used to predict on the equivalent set of bin pairs in HeLa-S3 and HepG2, which were subsequently reduced to those with enhancer targets only (using the same criteria as described above). Final probabilities were calculated based on the mean of probabilities over the 10 runs. The distance based F1-maximising cut-offs were applied to obtain a final list of interactions. Enhancer-promoter interaction sharing between GM12878, HeLa-S3 and HepG2 was calculated based on whether the interactions were present in 1, 2 or 3 of the cell types and venn diagrams were generated using the *R* package VennDiagram (Chen, 2016).

#### Example domains predicted in HeLa-S3 but not GM12878

Processed PolII ChIA-PET interactions data for HeLa-S3 was downloaded from ENCODE (The ENCODE Project Consortium, 2012) and start and end regions were lifted over to hg38, removing those without a 1:1 correspondence between the two builds. Bait-target pairs for predicted enhancer-promoter interactions in HeLa-S3 were selected according to a probability of at least 0.6 in HeLa-S3 and a probability of less than 0.4 in GM12878. Both ends were intersected with the lifted over start and end regions in the ChIA-PET data in order to generate a list of candidate examples. We selected an example on chromosome 6 due that chromosomes high performance statistics from the modelling described above. The *R* package Sushi (Phanstiel, 2015) was used to generate plots of the resulting loops and lines together with annotations and enhancer locations for the data sets in the example region.

### Analysis and predictions across 76 cell types from FANTOM5

#### Library selection and tanscriptional component generation

A total of 249 CAGE libraries FANTOM5 were selected according to the availability of sample replicates. Transcriptional decomposition was applied to generate PD and PI components for a total of 76 cell types (including 4 ENCODE cell lines and 72 primary cells - see Table S10 for list of library identifiers and names). For the purposes of consistency between the ENCODE generated data sets and the primary cell type generated data sets, the hyperparameters were fixed for the random walk and independent components to the same values which were generated from the models in ENCODE (whilst allowing the hyperparameters for the zero inflated negative binomial distribution to vary).

#### Hierarchical clustering of raw data samples

Raw binned data at 10kb resolution was normalised into tags per million (tpm) and the mean was taken across replicates to obtain a matrix of 76 columns against the total genomic bin count (303,065). Only regions potentially transcribed in the given set of CAGE assays were considered by asking for bins which had more than one cell type containing tags. The matrix was transformed into *log_10_* values (adding a pseudo-count of 1) and hierarchical clustering using `hclust` was applied in *R*. The ordering from the clustering was used to guide the row and column ordering in the heatmaps for Figure 5. The function cutree was used to find 10 groups from the clustering, which were annotated and merged manually to generate the most biologically relevant cell type groupings from the data.

#### Heatmaps for comparison PD and PI components

A common set of bins (34,953) was derived for comparison between cell types of the PD and the PI components by choosing the set of bins where the sign of the PI component was positive in more than one cell type. To generate similarity matrices, the PD and PI signals were converted into binary according to whether the signal was positive (1) or negative (0) (PI - 0.1 was used for the independent component to avoid non-expressed bins with very small positive estimates). The resultant matrix of 76 columns and rows according to the common bin set was used to calculate 1 - *L*_1_-norm between each pair of cell types to calculate a similarity matrix. The same metric to cell type boundary sharing and cell type enhancer promoter sharing, with methods described in the sections below. The *R* package `pheatmap` (Kolde, 2015) was used to generate cell type group annotated heatmaps of the similarity matrices.

#### Extending boundary predictions for 76 cell types from FANTOM5

For all cell types, XAD boundaries were calculated from the algorithm described above for ENCODE datasets, supplying the stability scores across the full set of cell types. To calculate boundary sharing across all cell types, non-overlapping bins of size of 100kb were generated and calculated for each expanded bin the number of cell types within which a boundary was found. In order to cover all possible windows the expanded bins were shifted by a 10kb bin 10 times and the final number of shared boundaries was calculated according to an average of the 10.

#### Predicting interactions for 76 cell types from FANTOM5

Datasets with features were generated as described for the ENCODE cell lines for all cell types, with the bin selection also extended to the full set, thus creating a common bin set which is larger than that for the ENCODE cell lines alone. Features non-specific to a cell type were also calculated more broadly with consideration to the 76 cell types. The model was trained as above using the GM12878 dataset using the broader bin set. Similar model performance was noted or this model when testing on the raw dataset using 10-fold cross validation. The held out set predicted probabilities were used for GM12878 and the predicted probabilities for the other 75 cell types were generated by averaging over 10 trained models across the whole dataset (to robustly account for random differences in the data balancing for the training).

#### Analysis of interaction data across samples

The datasets for the 76 cell types were reduced according to whether at least one dataset had an active CAGE enhancer annotated to it, in order to obtain a list of potential enhancer promoter interactions. To generate lists of predicted interactions, we generated distance based probability cut-offs in GM12878, using the same method for the ENCODE data sets above, using score cut-offs ≥ 3 and ≥ 5.

To calculate enhancer-promoter interaction sharing statistics, the ≥ 3 score cut off was used and for each possible number of cell types (from 1 to 76) the number of significant interactions which were present in exactly that number of cell types was calculated. To generate the enhancer-promoter sharing heatmap, all interactions which were present in more than one cell type were considered and pairwise cell type similarity calculated from (1 - L_1_-norm) between columns of the binary (1 predicted / 0 not predicted) matrix with cell types as columns and interaction set as rows. This generated a similarity matrix which we plotted using `pheatmap` (Kolde, 2015) based on the cell type ordering from the hierarchical clustering in the raw data.

#### Blood specific EP interactions

To isolate examples of interactions specific to blood, interactions meeting the criteria of present (≥ 3 score distance based efficient cut offs) in at least half of the cell types labelled as ‘blood cells’ (see Supplementary Table S11) and in fewer than a total of up to a maximum 3 more than the number of blood cells within the 76 cell types were classified as ‘blood specific’. For further trimming of examples and plotting of loops, probabilities assigned to all cell types assigned as blood cells were averaged into one using the mean. Merged probability vectors were also generated for endothelial cells, epithelial cells, muscle cells, mesenchymal and fibroblasts. Examples of blood specific interactions were selected based on the loop being still significant according to the averaged probabilities. The *R* package `Sushi` (Phanstiel, 2015) was used to generate plots of selected examples, including tracks for annotations and the averaged PD, PI components for blood cells.

#### GWAS enrichments across different components

The GWAS catalogue (Welter et al., 2014) for trait associated SNPs including SNPs in LD was downloaded for hg19 and coordinates lifted to hg38. For each trait with at least 50 assigned SNPs, a *χ*^2^ test was constructed per each of the 76 cell types and components (PD, PI, XAD boundaries EP interactions). For the PD and PI components, the foreground was assigned as the set of bins with a positive component sign in the given cell type, and the non-foreground assigned as the set of bins with a positive component sign in at least one cell type, but not the given cell type. For the EP interactions, the foreground was assigned as the set of target enhancers within the given cell type and the non-foreground based on the target enhancers within any other cell type. For the XAD boundaries, the foreground was assigned as the boundary bins +/− 2 bins in the given cell type against a non-foreground of all boundary bins +/− 2 bins in any cell type, not including the given cell type. The odds ratios was calculated based 2×2 tables including the number of trait SNPs within the foreground/non-foreground and the number of SNPs for all other trait SNPs within the foreground/non-foreground. For each component, the p-values from the total set of *χ*^2^ tests were corrected to an FDR with the Benjamini & Hochberg correction.

The library `pheatmap` (Kolde, 2015) was used to generate heatmaps of the resulting trait enrichments and the library `Sushi` (Phanstiel, 2015) was used to generate interaction plots for given EP interaction examples. To generate the heatmaps for the number of cell types with significant trait enrichments, the number of cell types with fdr < 0.01 and an odds of > 1.25 were counted per trait, per component. To calculate the component preference by trait, the resulting data was normalised per trait before generating the heatmap.

#### Resource generation of 76 cell types from FANTOM5

Values for the PD, PI components, together with locations of XAD boundaries and predicted enhancer-promoter interactions were saved out as BED files, using UCSC zero-based coordinates, and supplied as supplementary data files.

---

**Figure S1.**
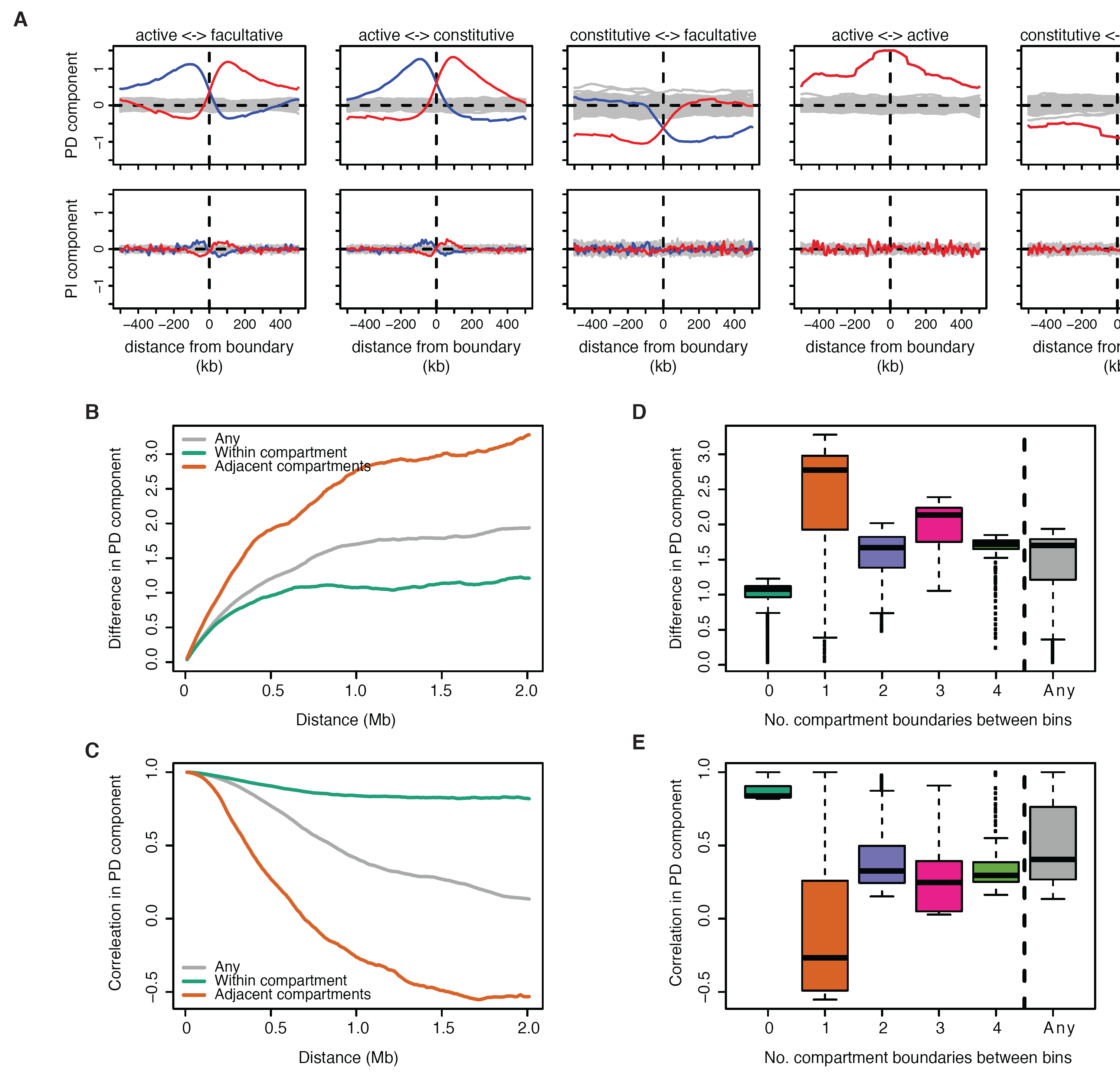
Relationship between transcriptional components and compartments. **A:** Average PD signal (top row) and PI signal (bottom row) around boundaries of HiC-derived chromatin compartments. Blue represents shifts from 5’ to 3’ genomic coordinates red represents shifts from 3’ to 5’. Horizontal dotted lines represent the transition between positive and negative and grey bands represent equivalent shifts across random compartment boundaries. **B-C:** Difference in PD component (**B**) and correlation (C) between cell lines, plotted according to distance between bins, for bin pairs falling within the same compartment (green), across adjacent compartments (orange) and any (grey). **D-E:** Box-and-whisker plots representing the distribution of differences in PD component (**D**) and correlations (**E**) across ENCODE cell lines, according to the number of compartments spanned by bin-pairs (up to 4 apart, and any).

**Figure S2.**
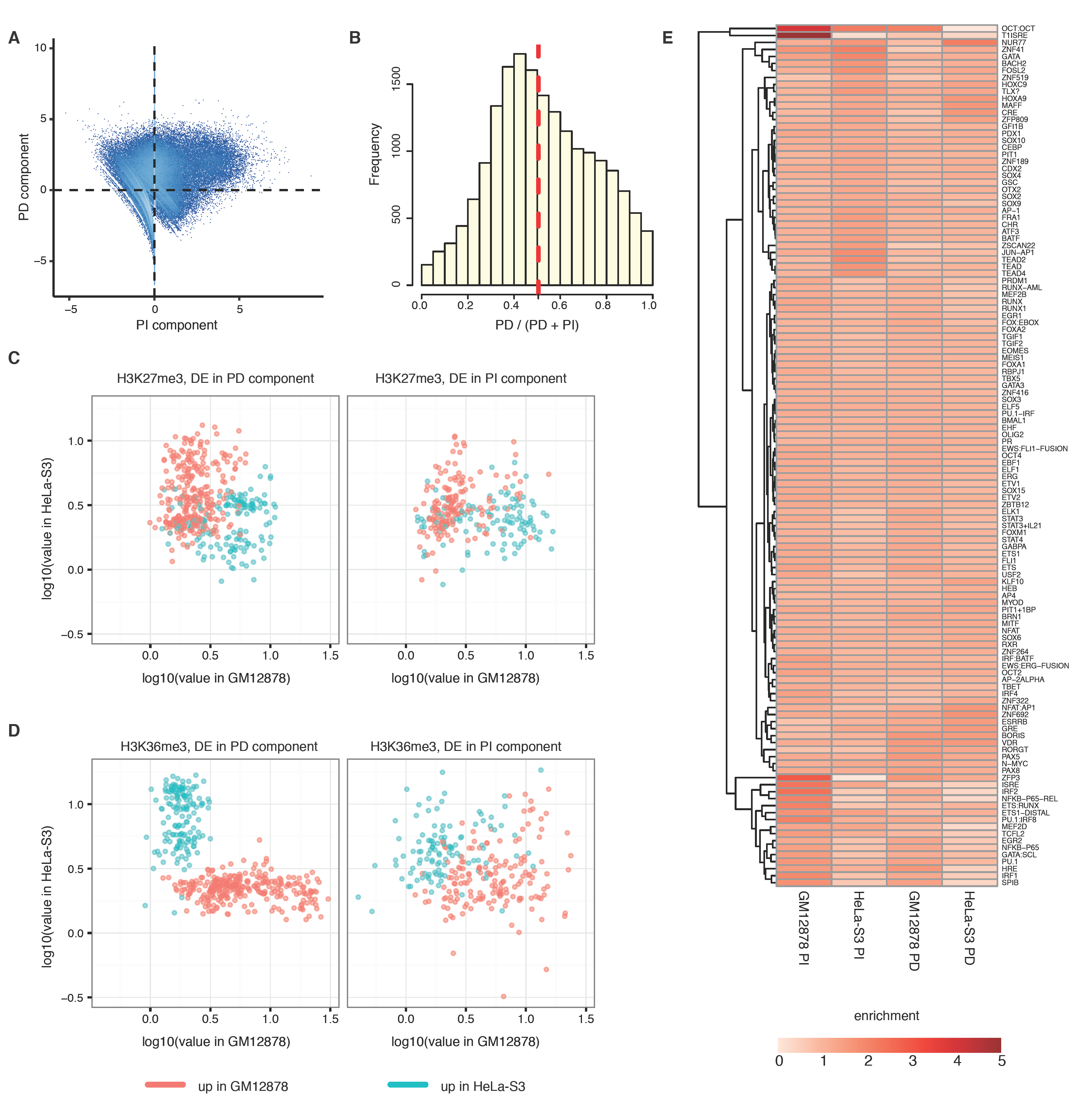
Separability of PD and PI components. **A**: Scatter plot of the PD component against the PI component in GM12878. Dotted lines represent the points where the components are equal to zero. **B**: Histogram of the ratio of the PD component to the sum of the PD and PI components, for bins containing expressed TUs. The frequency represents the number of TUs and the red dashed line represents the median. **C**: Differences in H3K27me3 levels (TPM normalised and aggregated bin-wise) between HeLa-S3 (vertical axes) and GM12878 (horisontal axes) cells at differentially expressed (DE) bins in the PD (left) and PI (right) components up in GM12878 (red) or up in HeLa-S3 (green). **D**: Differences in H3K36me3 levels (TPM normalised and aggregated bin-wise) between HeLa-S3 (vertical axes) and GM12878 (horisontal axes) cells at DE bins in the PD (left) and PI (right) components up in GM12878 (red) or up in HeLa-S3 (green).

**Figure S3.**
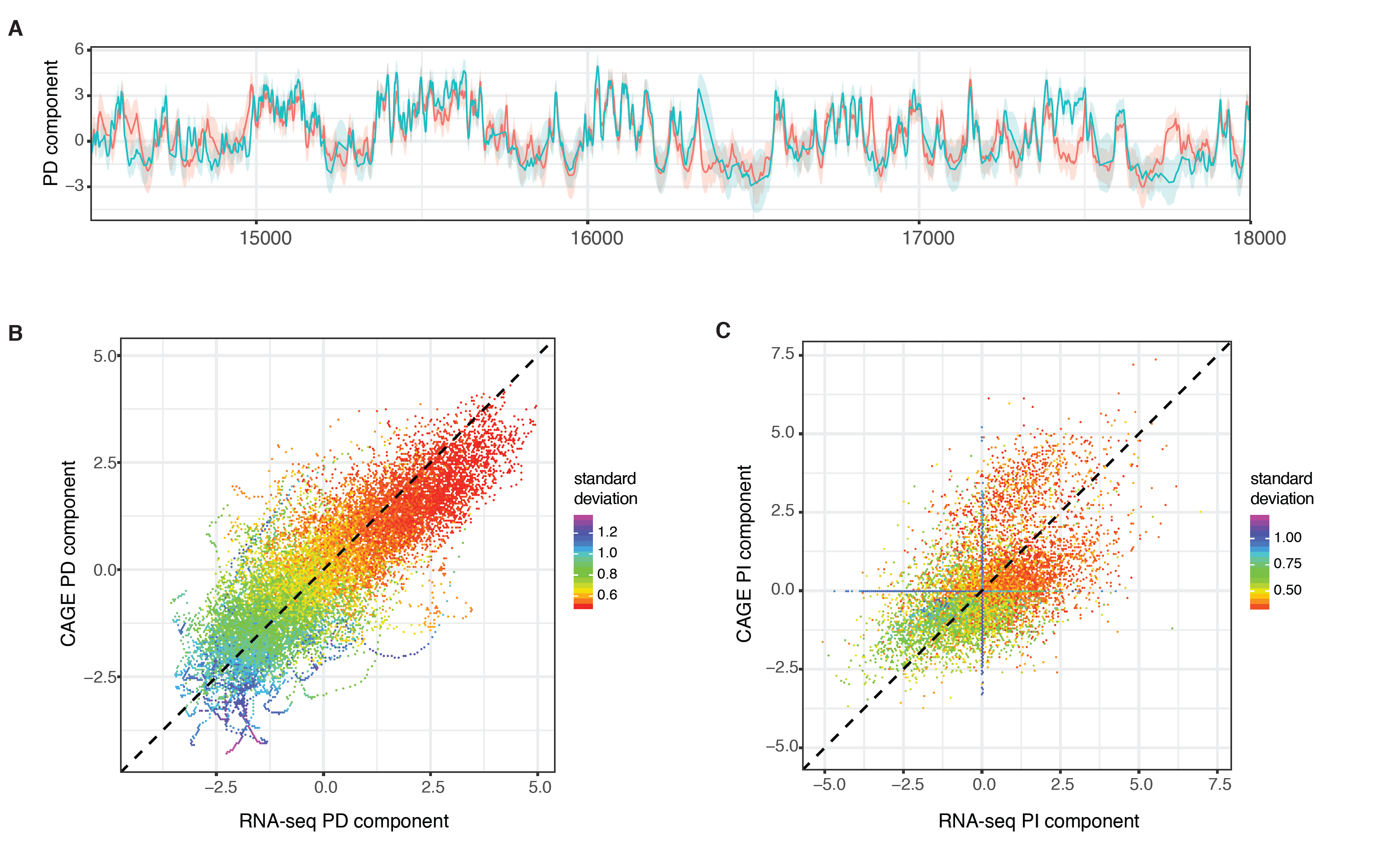
Transcriptional decomposition performed using RNA-seq data. **A**: PD components in GM12878 for CAGE (red) and RNA-seq (green), plotted (+/-) their respective standard deviations around their estimated posterior values. Locus plotted is chr1:145,000,000-180,000,000. **B-C**: Scatter plots of the PD component (**B**) and the PI component (**C**) for CAGE against RNA-seq, for all 10kb bins on chromosome 1 within 25 bins of an active TU. Colour scheme is generated according to the sums of the standard deviations from the two datasets.

**Figure S4.**
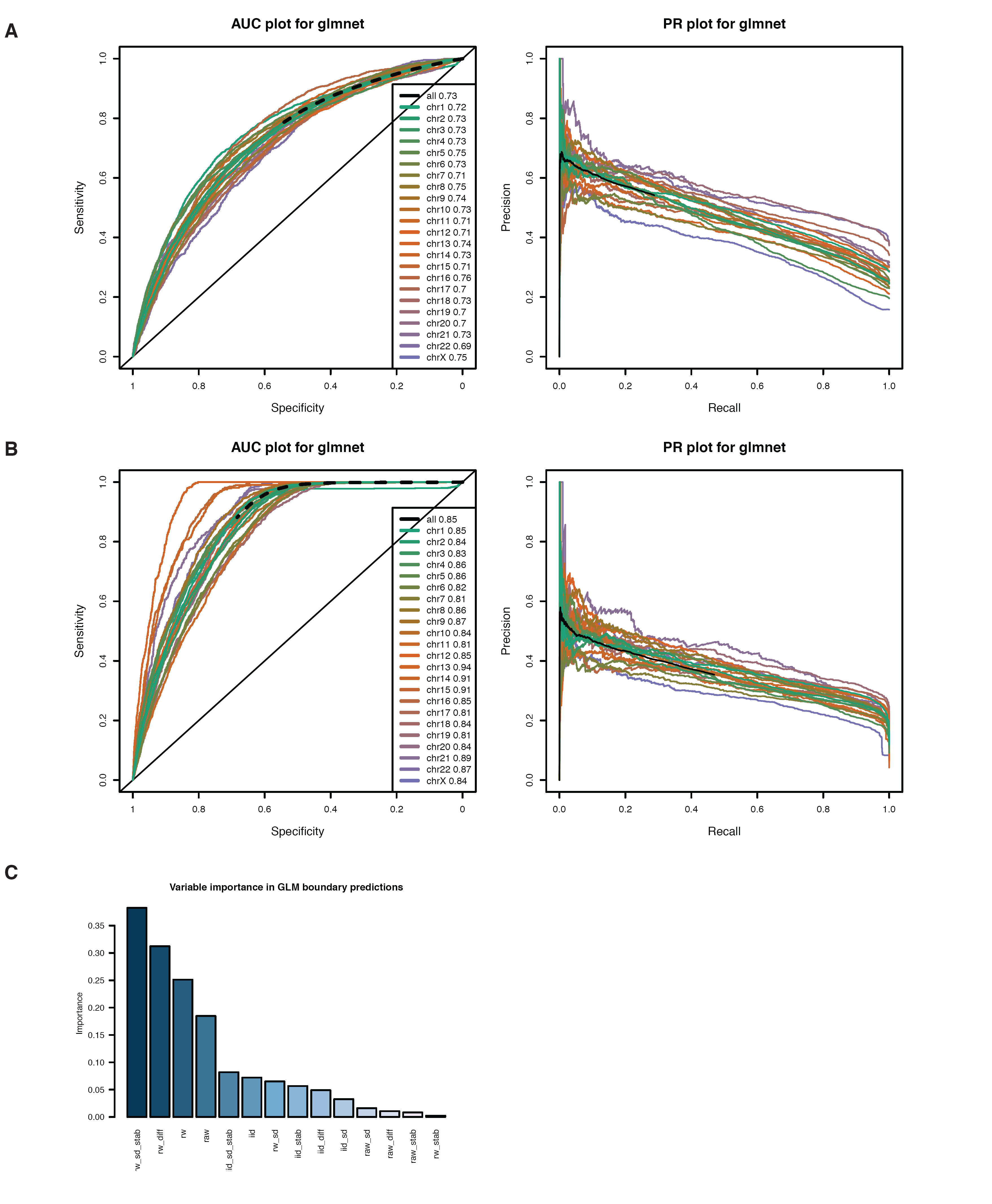
Predictability of TAD boundary regions from transcriptional components. **A-B**: ROC and PR for glmnet model, predicting all TAD boundary regions (**A**) and restricting on TAD boundary regions within positive PD regions (**B**). Performance based on held out data based on 2-fold cross validation and generated for the whole dataset as well as per chromosome. **C**: Relative importance of variables from the generalised linear model fit using the glmnet package.

**Figure S5.**
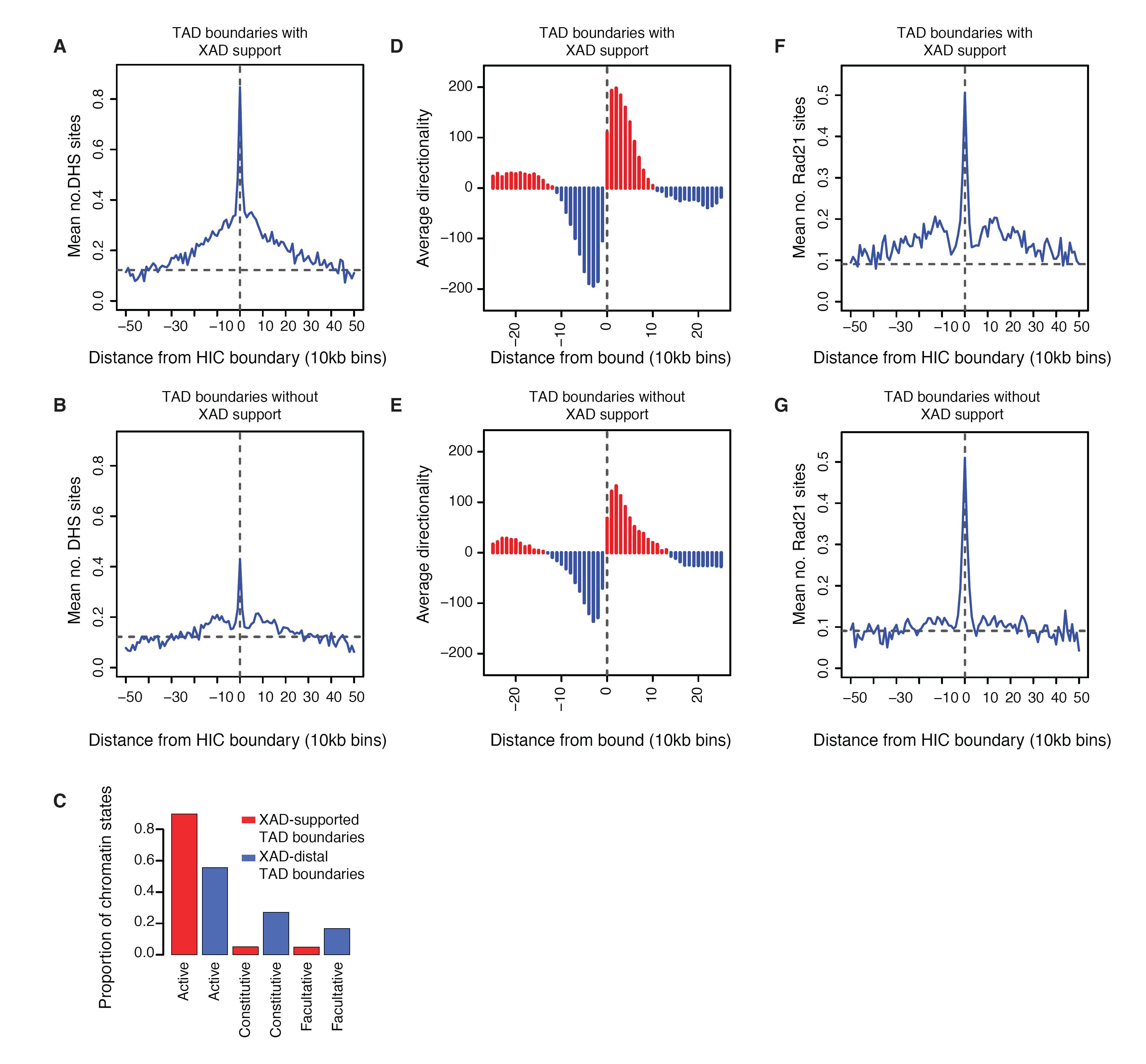
Properties of TAD boundaries according to XAD boundary support. **A-B**: Enrichment of DHS sites around TAD boundaries when supported by XAD boundaries (within +/-5 bins) (**A**) and when unsupported by XAD boundaries (**B**). **C**: Proportion of TAD boundaries within annotated chromatin environments, according to XAD boundary support. **D-E**: HiC directionality score around TAD boundaries when supported by XAD boundaries (within +/-5 bins) (**D**) and when unsupported by XAD boundaries (**E**). **F-G**: Enrichment of Rad21 sites around TAD boundaries when supported by XAD boundaries (within +/-5 bins) (F) and when unsupported by XAD boundaries (**G**).

**Figure S6.**
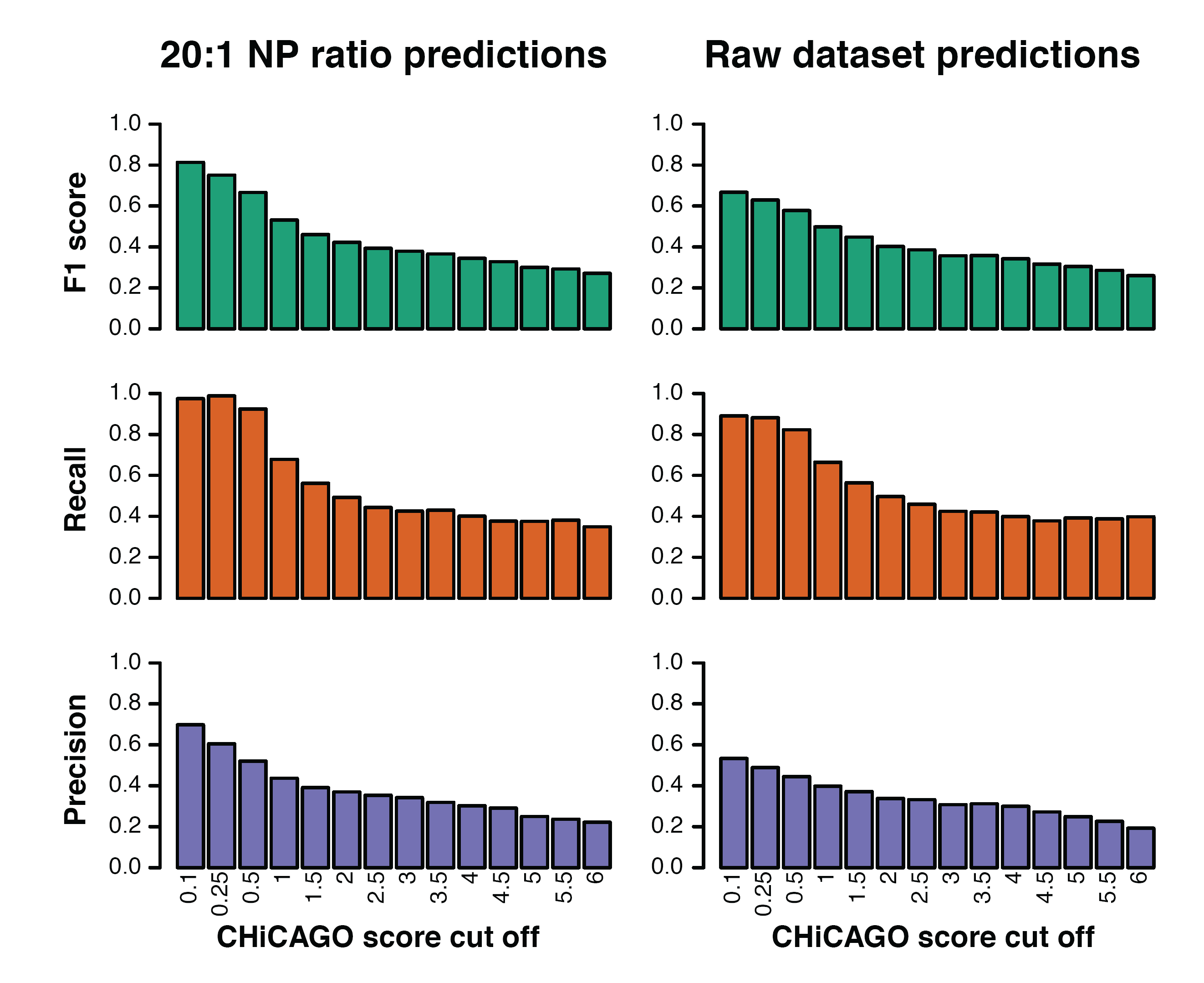
Effect of CHiCAGO score threshold on EP interaction prediction performance. F1 score, precision and recall of models for EP interactions, according to different CHiCAGO scores. All are based on training on a CHiCAGO score cut off of three and probability cut offs are based on F1 efficient cut-offs (distance variant) for each performance score threshold.

**Figure S7.**
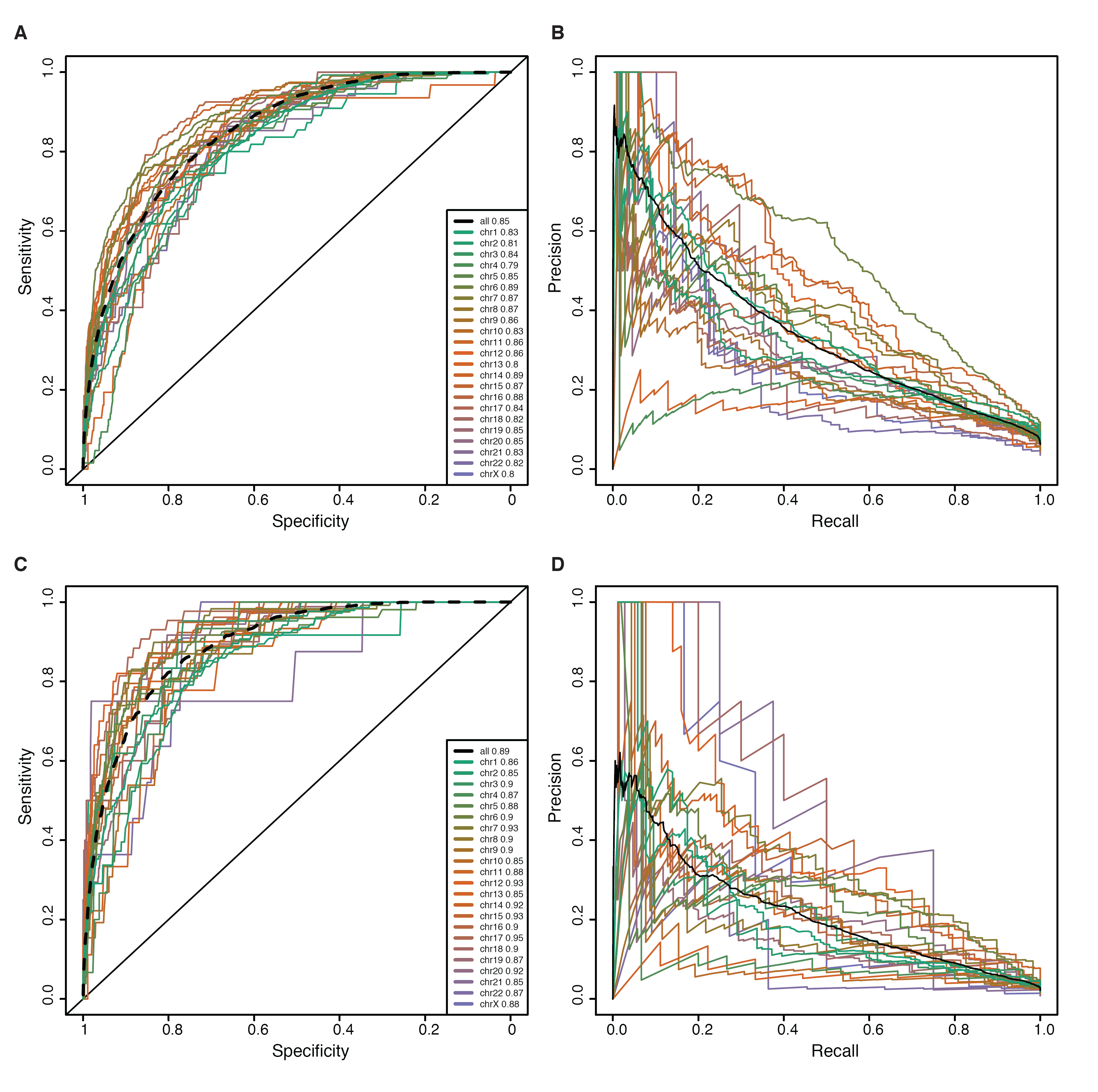
Global model performance for EP interactions, per chromosome. **A-D**: Chromosome specific ROC plots (**A,C**) and Precision-Recall plots (**B,D**) for performance in predicting EP interactions in GM12878, for CHiCAGO score ≥ 3 (**A,B**), or CHiCAGO score ≥ 5 (**C,D**). Black dashed lines represents performance across all chromosomes together.

**Figure S8.**
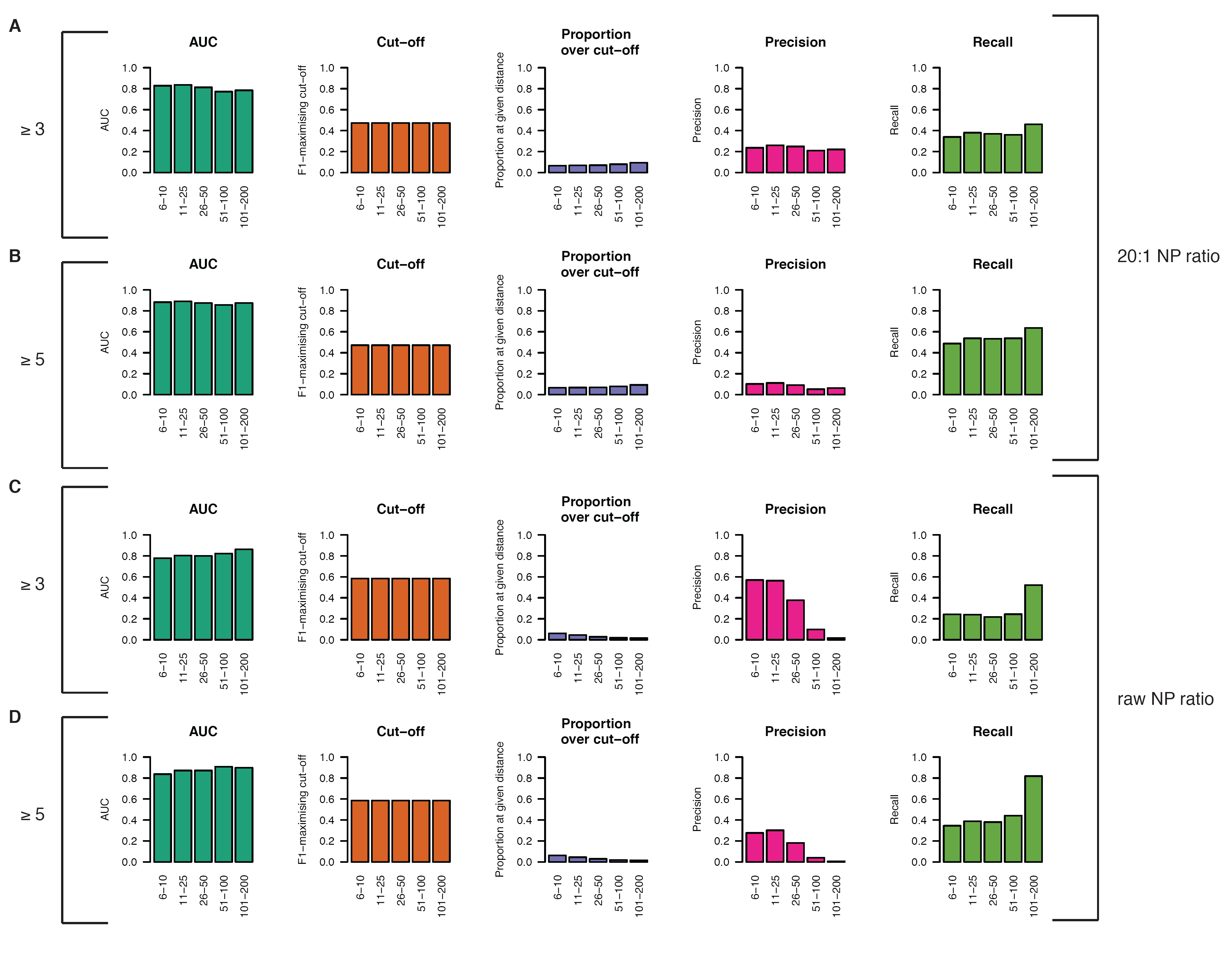
Model performance for EP interactions for a fixed cut-off over distance in GM12878. Fixed probability cut-off based on maximising the F1 statistic and performance statistics supplied separately over 5 distance groups. Results given for both the 20:1 NP balanced test dataset (**A,B**) and the raw test dataset (**C,D**), and according to CHiCAGO score ≥ 3 (**A,C**) and CHiCAGO score ≥ 5 (**B,D**).

**Figure S9.**
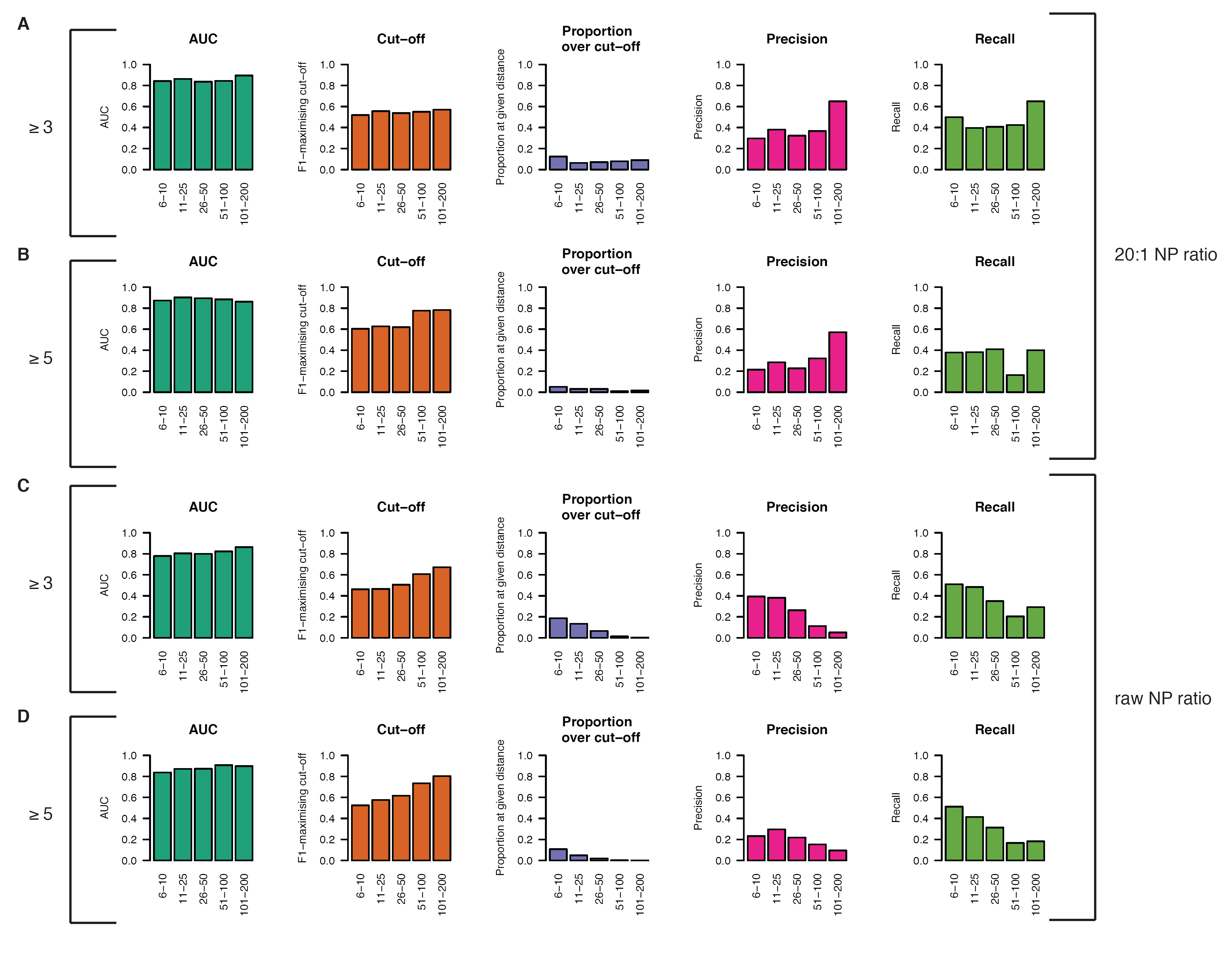
Model performance for EP interactions for varying cut-offs over distance in GM12878. Data split over 5 distance thresholds and cut-off chosen to be the F1 maximising cut-off at each threshold. Results given for both the 20:1 NP balanced test dataset (**A,B**) and the raw test dataset (**C,D**), and according to CHiCAGO score ≥ 3 (**A,C**) and CHiCAGO score ≥ 5 (**B,D**).

**Figure S10.**
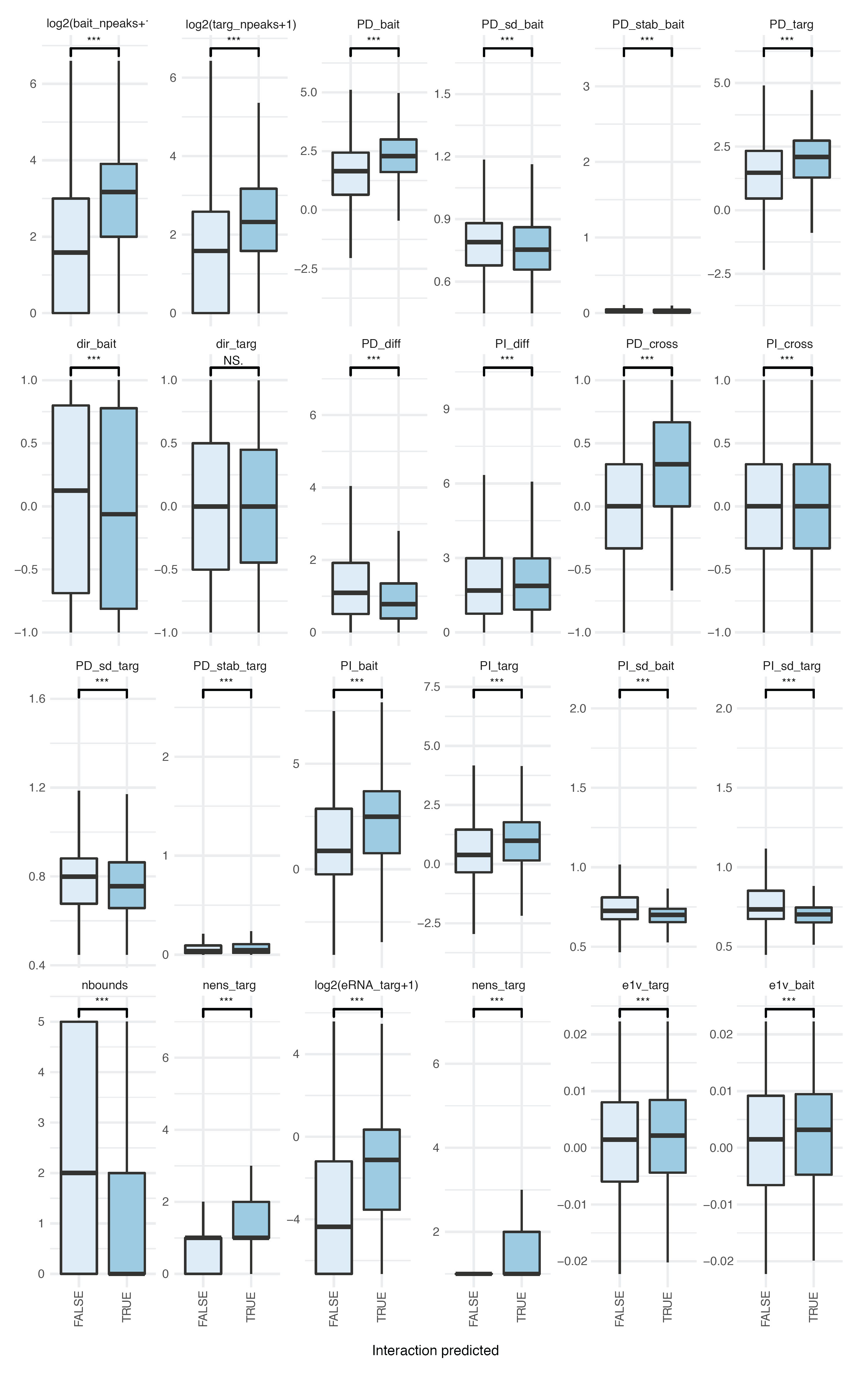
Features separating interacting vs non-interacting predicted enhancer-promoter interactions. Box and whisker plots for transcriptional based features, split according to predicted (darker blue) vs non-predicted (light blue) enhancer promoter interactions in GM12878. Significance based on pair-wise t-tests s.t. * = FDR<0.05, ** = FDR<0.01, *** = FDR<0.001 and NS = non-significant. Outliers not included in the box and whisker plots.

**Figure S11.**
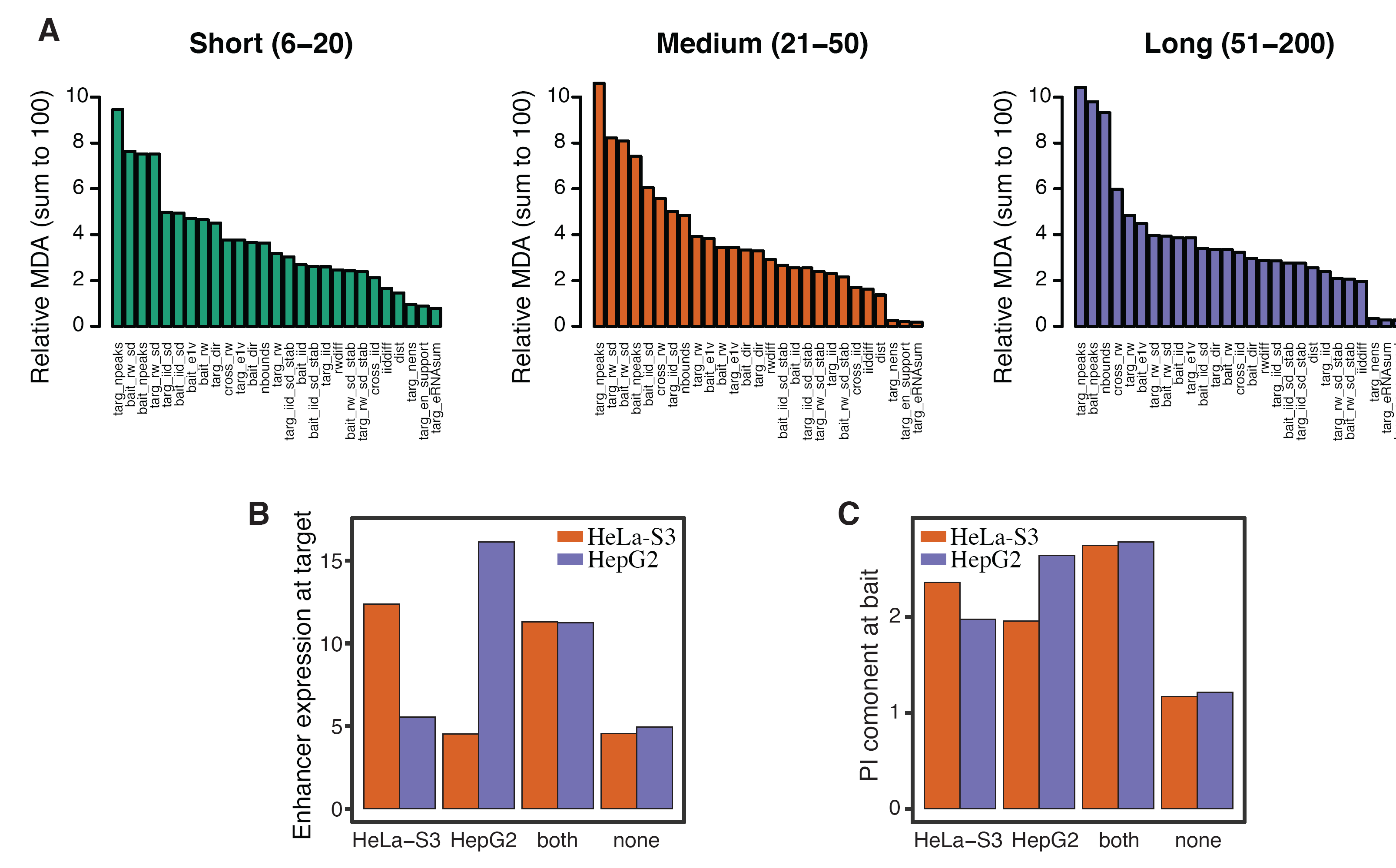
Feature importances for predicting enhancer-promoter interactions. **A**: Feature importance across three different sets of distances. **B-C**: Enhancer transcription at the target (**B**) and PI component at the bait (**C**) is enriched at cell-type specific predicted promoter-enhancer interactions. Bars represent mean feature levels for HeLa-S3 and HepG2 according to whether the interaction was predicted in HeLa-S3 only, HepG2 only, both, or none.

**Figure S12.**
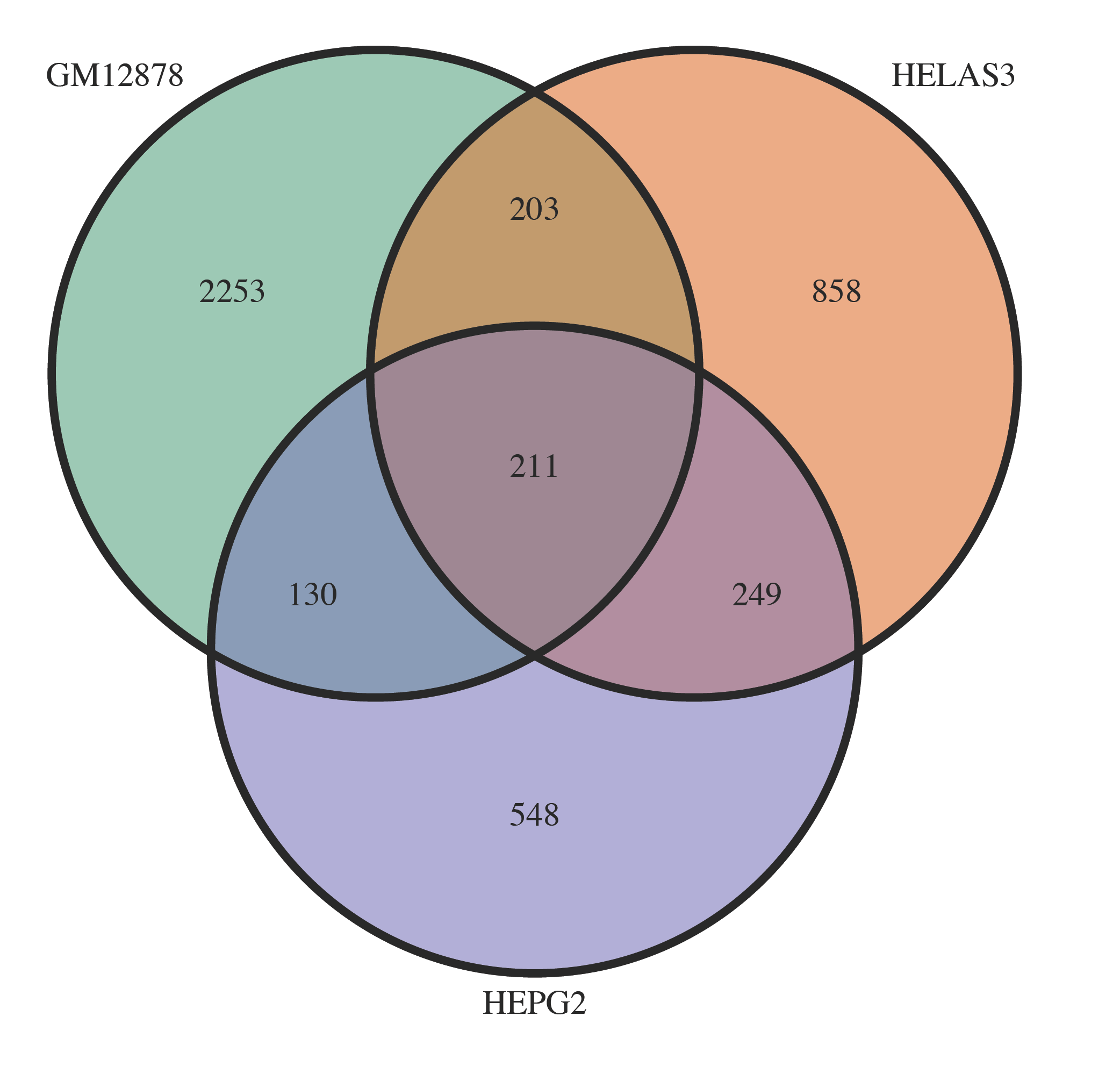
Effect of CHiCAGO score threshold on cell type differences in predicted EP interactions. Number of EP interactions shared in GM12878, HeLa-S3 and HepG2, for CHiCAGO score ≥ 5. Prediction cut off set by maximising the F1 statistic (distance variant) in GM12878.

**Figure S13.**
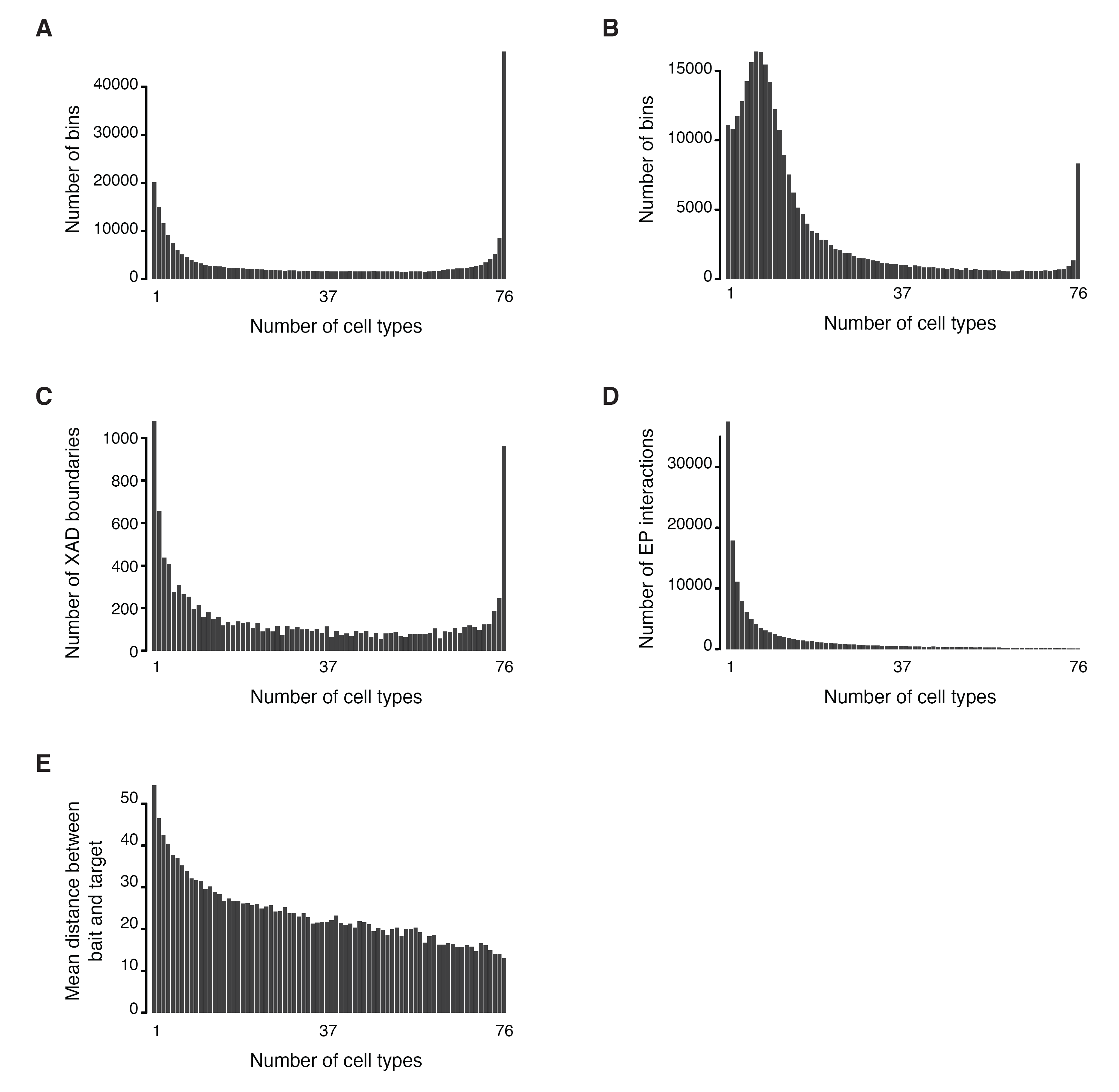
Similarities between cell types in transcriptional components. **A**: Sharing of bins with positive PD signal across cell types. **B**: Sharing of bins with positive PI signal across cell types. **C**: Sharing of XAD boundary regions across cell types at 100kb resolution. **D**: Sharing of predicted EP interactions across cell types. For all interactions present in at least one cell type, bars represent the number of cell types in which it was predicted. **E**: Sharing of predicted EP interactions across cell types versus mean distance between bait (promoter) and target (enhancer).

**Figure S14.**
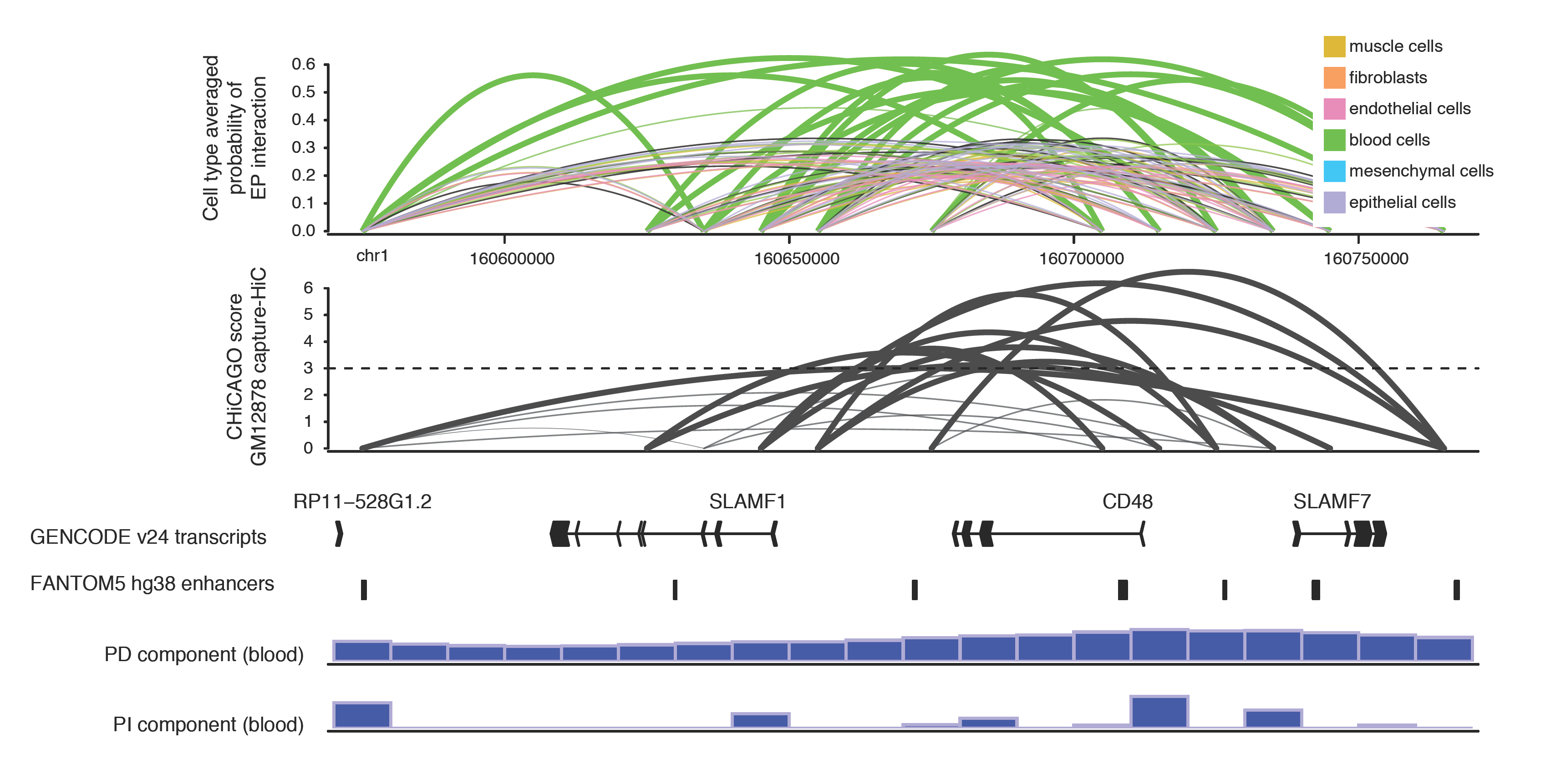
Transcriptional decomposition reveals cell type specific regulatory organisation around *CD48* gene. From top to bottom: predicted probability of EP interactions averaged across groups of cells (as in Figure 5), CHiCAGO score of interaction based on GM12878 capture HiC data. Below are displayed (in the following order) GENCODE v24 transcripts, FANTOM5 enhancers, the average PD and PI component across blood cells.

**Figure S15.**
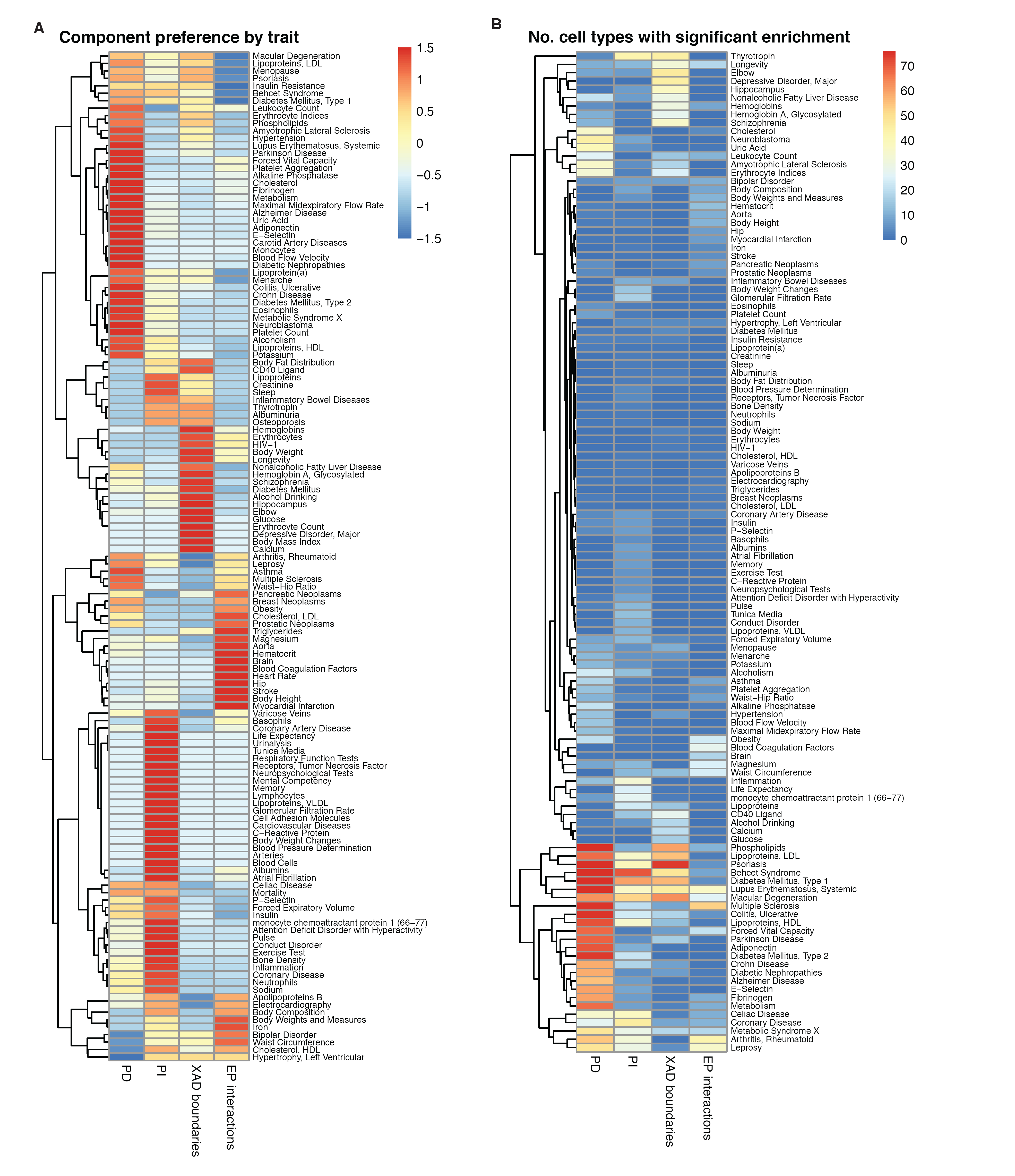
Component and cell type preferences of GWAS trait SNP enrichments. **A**: Preferences of GWAS trait-associated SNPs in PD positive bins, PI positive bins, XAD boundaries and EP interactions identified in 76 human cell types. Preference calculated based on the number of significant (FDR<0.01 and odds>1.25) cell types per trait and component (see methods), scaled across the four components. **B**: Number of cell types with significant (as in **A**) enrichments of GWAS trait-associated SNPs in PD positive bins, PI positive bins, XAD boundaries and EP interactions identified in 76 human cell types.

**Figure S16.**
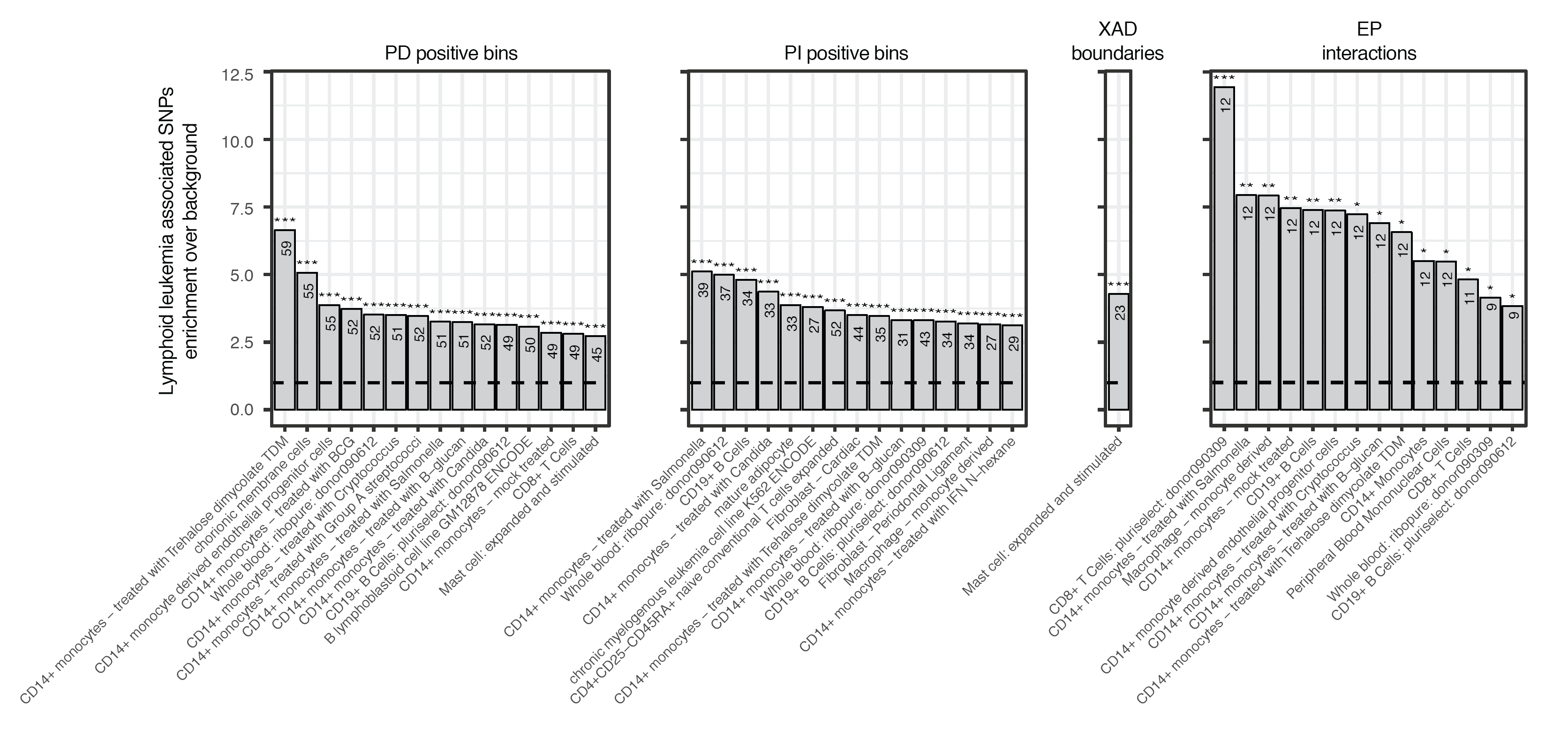
Analysis of lymphoid leukemia associated SNPs reveals cell-type and expression component preferences. Lymphoid leukemia SNP enrichments in transcriptional components, XAD boundaries, and inferred enhancer-promoter interactions (FDR-corrected *χ*^2^ tests based on the PD/PI +ve bins or presence of XAD boundaries/target enhancers per cell type, and trait associated SNPs from the GWAS catalog, see methods)

**Table S1.** Best correlation parameter combinations for RNA-seq transcriptional decomposition, per chromosome.

| Chromosome | size | prob | RW | IID | PD correlation | PI correlation |
| --- | --- | --- | --- | --- | --- | --- |
| 1 | 8 | -1 | -4 | 2 | 0.8343338 | 0.8685478 |
| 2 | 8 | -1 | -7 | -1 | 0.8314016 | 0.8384131 |
| 3 | 8 | -4 | -4 | 2 | 0.8426355 | 0.8464501 |
| 4 | 8 | 3 | -4 | 2 | 0.8047164 | 0.7977146 |
| 5 | 8 | -4 | -4 | 2 | 0.8168530 | 0.8325121 |
| 6 | 8 | -4 | -4 | 2 | 0.8068548 | 0.8185471 |
| 7 | 8 | -4 | -6 | -1 | 0.8285529 | 0.8204263 |
| 8 | 8 | -4 | -4 | 2 | 0.8088375 | 0.8117918 |
| 9 | 2 | 3 | -4 | 2 | 0.8329053 | 0.9064663 |
| 10 | 7 | -4 | -4 | 2 | 0.8109703 | 0.8298349 |
| 11 | 2 | -4 | -4 | 2 | 0.8575597 | 0.8587555 |
| 12 | 2 | -4 | -4 | 2 | 0.8514417 | 0.8623694 |
| 13 | 7 | 3 | -3 | 2 | 0.8114826 | 0.9815640 |
| 14 | 8 | 2 | -3 | 2 | 0.7948931 | 0.9673565 |
| 15 | 8 | -4 | -3 | 2 | 0.8277752 | 0.9636634 |
| 16 | 2 | 2 | -3 | 2 | 0.8642273 | 0.8265529 |
| 17 | 2 | -4 | -3 | 2 | 0.8772700 | 0.8866936 |
| 18 | 1 | 3 | -6 | -1 | 0.7759199 | 0.7527304 |
| 19 | 2 | -4 | -3 | 2 | 0.8981290 | 0.9118324 |
| 20 | 7 | -4 | -3 | 2 | 0.8327296 | 0.8354974 |
| 21 | 1 | -3 | -5 | -1 | 0.7631566 | 0.9470045 |
| 22 | 7 | 2 | -5 | -1 | 0.8484782 | 0.9509241 |
| X | 7 | 3 | -3 | 2 | 0.8150549 | 0.8073811 |

**Table S2.** Differentially expressed bins in the PD component. Based on posterior estimates of the difference between bins in GM12878 VS HeLa-S3. Bins with FDR< 0.01, corrected according to the number of raw-expressed bins, are considered significant. Score listed as 1 if the bin PD component was significant up in HeLa-S3 vs GM12878, and -1 if up in GM12878 vs HeLa-S3.

**Table S3.** Differentially expressed bins in the PI component. Based on posterior estimates of the difference between bins in GM12878 VS HeLa-S3. Bins with FDR< 0.01, corrected according to the number of raw-expressed bins, are considered significant. Score listed as 1 if the bin PD component was significant up in HeLa-S3 vs GM12878, and -1 if up in GM12878 vs HeLa-S3.

**Table S4.** Features used for TAD GLM training. PI and PD features based on chromosomal transcriptional decomposition model for GM12878. Abbrev. gives the abbreviated names for the features and Description explains how the features were derived.

| Feature | Abbrev. | Description |
| --- | --- | --- |
| PD signal | PD | Posterior PD estimate |
| PD standard deviation | PD_sd | Posterior PD standard deviation |
| PD stability | PD_stab | Standard deviation across cell types of PD_sd |
| PD difference | PD_diff | First order difference of PD |
| PI signal | PI | Posterior PI estimate |
| PI standard deviation | PI_sd | Posterior PI standard deviation |
| PI stability | PI_stab | Standard deviation across cell types of PI_sd |
| PI difference | PI_diff | First order difference of PI |
| RAW signal | raw | Mean log<sub>2</sub> TPM over replicates |
| RAW difference | raw_diff | First order difference of raw |
| RAW standard deviation | raw_sd | Standard deviation of log<sub>2</sub> TPM over replicates |

**Table S5.** List of features trained against significant proximity interactions. Features calculated for the bait or/and the target separately, or using information from both. Features calculated based using information from all cell types are listed as non-cell type specific (N), in contrast to those which may be calculated only from the cell type of interest (Y). Data type listed as continuous (C) or discrete (D), whereby the numbers in brackets indicate the possible values which the feature may take.

| Term | Abbreviation(s) | Cell-type specific? | Data type |
| --- | --- | --- | --- |
| PD value at bait/target | PD_bait,PD_targ | Y | C |
| PD sd value at bait/target | PD_sd_bait PD_sd_targ | Y | C |
| PD stability at bait/target | PD_stab_bait,PD_stab_targ | N | C |
| PI value at bait/target | PI_bait,PI_targ | Y | C |
| PI sd value | PI_sd_bait,PI_sd_targ | Y | C |
| PD difference | PD_diff | Y | C |
| PI difference | PI_diff | Y | C |
| PD cross correlation | PD_cross | N | C |
| PI cross correlation | PI_cross | N | C |
| Enhancer expression at target | eRNA_targ | Y | C |
| Directionality at bait/target | dir_bait, dir_targ | Y | C |
| First eigenvector of PD cross correlation | e1v_bait, e1v_targ | N | C |
| XAD boundary insulation | nbounds | Y | D (0-3) |
| Distance between bins | dist | N | D (6-200) |
| Peaks detected at bait/target | npeaks_bait,npeaks_targ | Y | D (0+) |
| Predicted enhancers at target | nens_targ | Y | D (0+) |
| No. cell lines supporting target enhancer | en_support_targ | N | D (0+) |

**Table S6.**
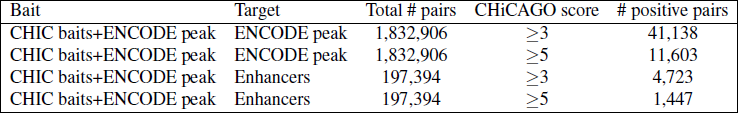
Number of positives called from the promoter-capture HiC GM12878 data, ENCODE cell line datasets set. ENCODE peak refers to a peak transcribed in more than one replicate in at least one of the ENCODE datasets. Enhancers refers to a CAGE defined enhancer which is transcribed in at least one of the ENCODE datasets.

| Bait | Target | Total # pairs | CHiCAGO score | # positive pairs |
| --- | --- | --- | --- | --- |
| CHIC baits+ENCODE peak | ENCODE peak | 1,832,906 | $\geq 3$ | 41,138 |
| CHIC baits+ENCODE peak | ENCODE peak | 1,832,906 | $\geq 5$ | 11,603 |
| CHIC baits+ENCODE peak | Enhancers | 197,394 | $\geq 3$ | 4,723 |
| CHIC baits+ENCODE peak | Enhancers | 197,394 | $\geq 5$ | 1,447 |

**Table S7.** Predicted GM12878 EP interactions. Bin set based on intra-chromosomal enhancers and baits active across ENCODE cell lines. Predictions derived from 10-fold validation testing on model training on GM12878 CaptureHIC CHICAGO score-derived significant interactions. EP interactions predicted for score cut-off ≥ 3 and ≥ 5. See methods for further details.

**Table S8.** Predicted HeLa-S3 EP interactions. Bin set based on intra-chromosomal enhancers and baits active across ENCODE cell lines. Predictions derived from testing HeLa-S3 features from model training on GM12878 CaptureHIC CHICAGO score-derived significant interactions. EP interactions predicted for most efficient cut-off based on score ≥ 3 and score ≥ 5. See methods for further details.

**Table S9.** Predicted HepG2 EP interactions. Bin set based on intra-chromosomal enhancers and baits active across ENCODE cell lines. Predictions derived from testing HepG2 features from model training on GM12878 CaptureHIC CHICAGO score-derived significant interactions. EP interactions predicted for most efficient cut-off based on score ≥ 3 and score ≥ 5. See methods for further details.

**Table S10.** Number of positives called from the promoter-capture HiC GM12878 data, 76 datasets set. Any peak refers to a peak transcribed in more than one replicate in at least one of the 76 datasets. Enhancers refers to a CAGE defined enhancer which is transcribed in at least one of the 76 datasets.

| Bait | Target | Total # pairs | CHiCAGO score | # positive pairs |
| --- | --- | --- | --- | --- |
| CHIC baits+Any peak | Any peak | 4,012,550 | $\geq 3$ | 71,044 |
| CHIC baits+Any peak | Any peak | 4,012,550 | $\geq 5$ | 18,495 |
| CHIC baits+Any peak | Enhancers | 863,039 | $\geq 3$ | 16,989 |
| CHIC baits+Any peak | Enhancers | 863,039 | $\geq 5$ | 4,617 |

**Table S11.**
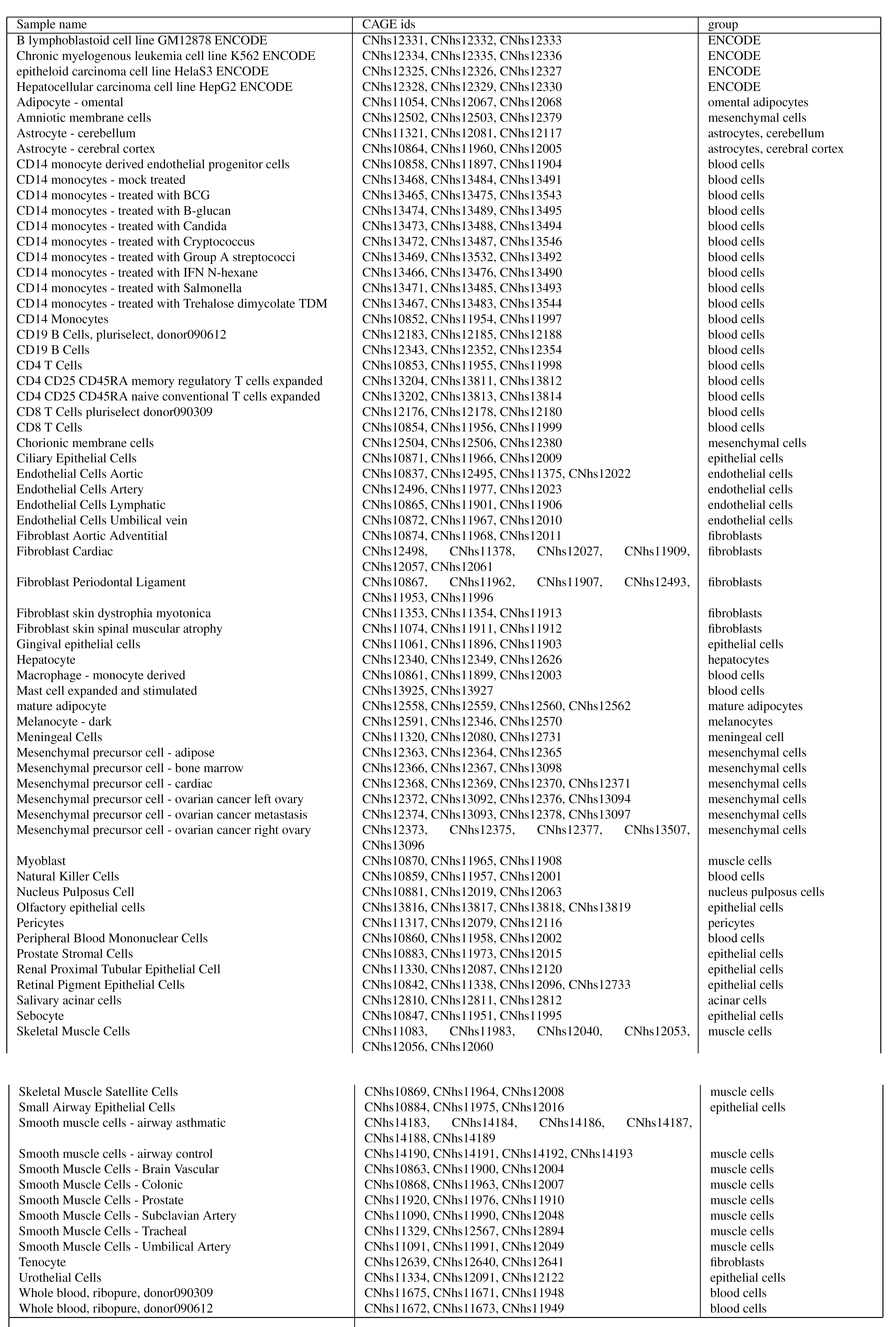
Replicated CAGE samples and associated FANTOM5 ids used in the analysis.

